# Advancements in Vaccine Development : A comprehensive design of a Multi-Epitopic Immunodominant Peptide Vaccine Targeting Kyasanur Forest Disease via Reverse Vaccinology

**DOI:** 10.1101/2025.02.03.634026

**Authors:** Sarika Baburajan Pillai, Akshay Jeyachandran, Naseera Kannanthodi Pariyapurath, Sumitha Jagadibabu, Paramasivan Rajaiah, Shivanandappa Kukkaler Channappa, Rahul Gandhi Pachamuthu, Ananda Arona Premkumar, Selvaraj Jagannathan

**Affiliations:** Pasteur Institute of India, Coonoor, The Nilgiris, Tamilnadu, India; University of Wuerzburg, Bavaria, Germany; Justice Basheer Ahmed Sayeed College for Women (Autonomous), Chennai, Tamilnadu, India; Division of Molecular Biology and Diagnostics, Vector Control Research Centre Field Station, ICMR, Madurai, Tamil Nadu, India; Centre for Nano Science and Technology, The Madanjeet School of Green Energy Technologies, Pondicherry University, Puducherry, India

**Keywords:** Kyasanur Forest Disease, Multi-epitope vaccine, *in silico*, Zoonosis, Immunoinformatic, Emerging Infectious Disease

## Abstract

Kyasanur Forest Disease (KFD), caused by the Kyasanur Forest Disease Virus (KFDV), is a tick-borne hemorrhagic fever virus first identified in Karnataka, South India, in 1957. The rising incidence of cases is alarming, particularly due to the potential spread of the disease to neighboring states. There presently exists no particular treatment, and the Chick embryo fibroblast (CEF) vaccine only provides temporary immunity and necessitates many doses, which results in poor immunisation rates among at-risk groups. This situation highlights the urgent need for a highly effective and user-friendly vaccine. Given that KFDV is classified as a BSL-4 pathogen, this study employs advanced immuno-informatics tools to design a safer vaccine construct. Promising results have identified the E protein of KFDV as an immunodominant target that can stimulate robust immune responses, including the production of neutralizing antibodies.

The vaccine design utilized various *in silico* tools to predict CTL, HTL and B-cell epitopes from the E protein, ensuring powerful immunological reactions. Quality attributes such as antigenicity, allergenicity, toxicity, and immunogenicity were assessed using specialized tools. A total of 16 epitopes (5 CTL, 3 HTL, and 8 B-cell) were identified, all exhibiting desirable properties like non-toxicity, non-allergenicity, high antigenic and immunogenic value. These epitopes were linked to form the PKFDVac-I vaccine, consisting of 279 amino acids with a MW of 29,162.32 Daltons. The constructed vaccine demonstrated stability and a robust immune response when docked with the TLR-4 receptor, indicating strong immunogenic potential. This multi-epitope peptide vaccine candidate developed via user-friendly and safer bio-informative tools represents a significant advancement in effective immunization strategies against KFDV infection, paving the way for future vaccine development efforts targeting other emerging infectious diseases. In conclusion, the integration of diverse epitopes into a cohesive vaccine prototype demonstrates a promising avenue for custom synthesis and application in immunization strategies. Further validation through *in vitro* and *in vivo* studies is essential to authorize the effectiveness of the designed vaccine construct.

## INTRODUCTION

Kyasanur Forest Disease is an arboviral illness that causes fever and bleeding, spread by arthropods, particularly ticks[1]. Viral diseases account for a considerable burden of global morbidity and mortality, with many being zoonotic in nature, originating in animal hosts before crossing species barriers to infect humans[2]. The zoonotic virus was found to have been distributed in Karnataka, South India (1957) [3]. After a 3–8 day incubation period, KFD symptoms which include chills, fever, and headaches appear abruptly. The manifestation of intense myalgia, emesis, gastrointestinal disturbances, and hemorrhagic complications may present three days subsequent to the initial onset of symptoms[4].Case studies on KFD indicated that it was restricted to the Karnataka’s arboraceous Shimoga district, recent reports indicate that it has spread to Tamil Nadu (Nilgiris), Kerala (Wayanad & Malappuram), Maharashtra (Sindhudurg), and Goa (Pali)[5]. From 1957 to 2017, approximately 9,594 cases of Kyasanur Forest Disease (KFD) have been reported across various districts along India’s western coast. Since 1957 till 2020, around 3,314 monkey deaths have been ascribed to KFD[6]. On an annual basis, approximately 160 instances of human cases are documented, exhibiting a CFR of 2.4% [7]. The disease is labeled as seasonal usually reported from December to May[8]. A study conducted on serosurvey samples from Kingaon-West Bengal (1962), Kutch and Saurashtra-Gujarat (1971), Parbatpur-Rajasthan (1971), and the Andaman and Nicobar Islands (2002) found evidence of hemagglutination inhibition antibodies of KFDV[9]. Chinese researchers have reported similar diseases caused by the Nanjianyin virus[10], and Saudi researchers have reported the Alkhurma Hemorrhagic Fever (AHF) since 1995[11] [12] in the dry season. Individuals who trekked to forests [9] for the purpose of collecting firewood, grasses, and various other forest-derived products were observed to exhibit instances of human cases [13]. The most typical way for ticks to spread the infection to people is by contact with an infected host, particularly one that is either ill or has recently succumbed to illness, such as a monkey[14]. Human-to-human transmission is not yet reported in the case of KFD[15].

A number of vaccines were tested to combat the disease in addition to the presently in use formalin-inactivated CEF KFD vaccine[16]. Studies carried out in KFD affected districts of Karnataka during 1990-92 reported an efficacy of 79.3% for one dose of the vaccine, and 93.5% for two doses and in 2005-10, effectiveness was found to be 62.4% in subjects who took two doses and 82.9% in those who took a booster dose after 2 doses [17], [18]. Some researchers have speculated that the lower efficacy of the vaccine may be due to drifts and diversification from KFDV strains currently circulating compared to the strain originally used to make the vaccine. This indicates a clear need for a detailed study and a highly effective vaccine to contain the spread of the disease[19].

Given that the KFD virus is classified within the Biosafety Level 4 (BSL-4) category of pathogens, this study has employed advanced methodologies such as promising reverse vaccine approach and immuno-informatic tools in an effort to design a multi-epitope peptide vaccine (MEPV) construct aimed at inducing an immune response. MEPV target specific epitopes of the pathogen and potentially improve efficacy. Vaccines that target multi epitopes are more successful as they lessen the possibility that the pathogen may evolve escape mutants. Since they don’t contain entire pathogens: live or attenuated, they are also generally safe. Furthermore, these vaccines are very adaptable, enabling customization to meet the demands of specific individuals or populations by taking genetic variability into account.

The diameter of KFD virus is about 25 nm and the length of the +ve strand RNA genome is nearly 11 bp[20]. It encodes a single polyprotein comprising of the structural proteins - E, C and prM/M and the non-structural proteins (N S-1, N S-2 a&b, N S-3, N S-4 a&b, and N S-5) after translation[21]. The C protein of KFDV plays two crucial roles in the infection process. It encapsulates viral RNA for protection, and also interacts with various host proteins to stimulate virus replication. Nonstructural proteins are proteases that are primarily responsible for cleaving the polyproteins and regulating host cell response either by directly interacting with the C protein, NS-2a and NS-3 aid in virus assembly or through interactions with structural proteins, NS1 regulates the production of infectious particles[22].

Immunodominant protein E of KFDV, like other flavivirus E proteins, aids in receptor binding and entrance into host cells via the fusing of the viral and cellular membranes[23]. There exists evidence suggesting that within the structural proteins of KFDV and other flaviviruses, the E protein is believed to have an imperative role in the virus’s virulence or is essential in determining tissue tropism [24]. When a virus invades a host, it attaches to the cell’s surface with the help of the E-protein, and the neutralizing antibodies produced in the second week recognize epitopes mostly found in the E-glycoprotein of flaviviruses[25]. Furthermore, computational studies have illustrated that the E protein exhibits significant immunodominance, heightened antigenicity, and a reduced propensity for eliciting allergic responses when compared to other proteins associated with KFD, thereby rendering it an optimal candidate for the advancement of innovative vaccines [26] and methodologies[27].

## MATERIALS AND METHODS

### Selection of KFD Strain to Retrieve the Protein Sequence

To identify a suitable strain for this study, KFD Virus species isolated and gene sequenced between the years 1957 to 2020 was obtained from NCBI and analyzed. The data revealed that the strains were isolated and genes were sequenced from humans, natural hosts, and reservoirs of KFD infection and also the vector (tick -*Haemaphysalis spp*.,)[28]. Table 1 shows the details of the sequences of KFDV isolated from humans and shortlisted for this study.

**Table 1:**
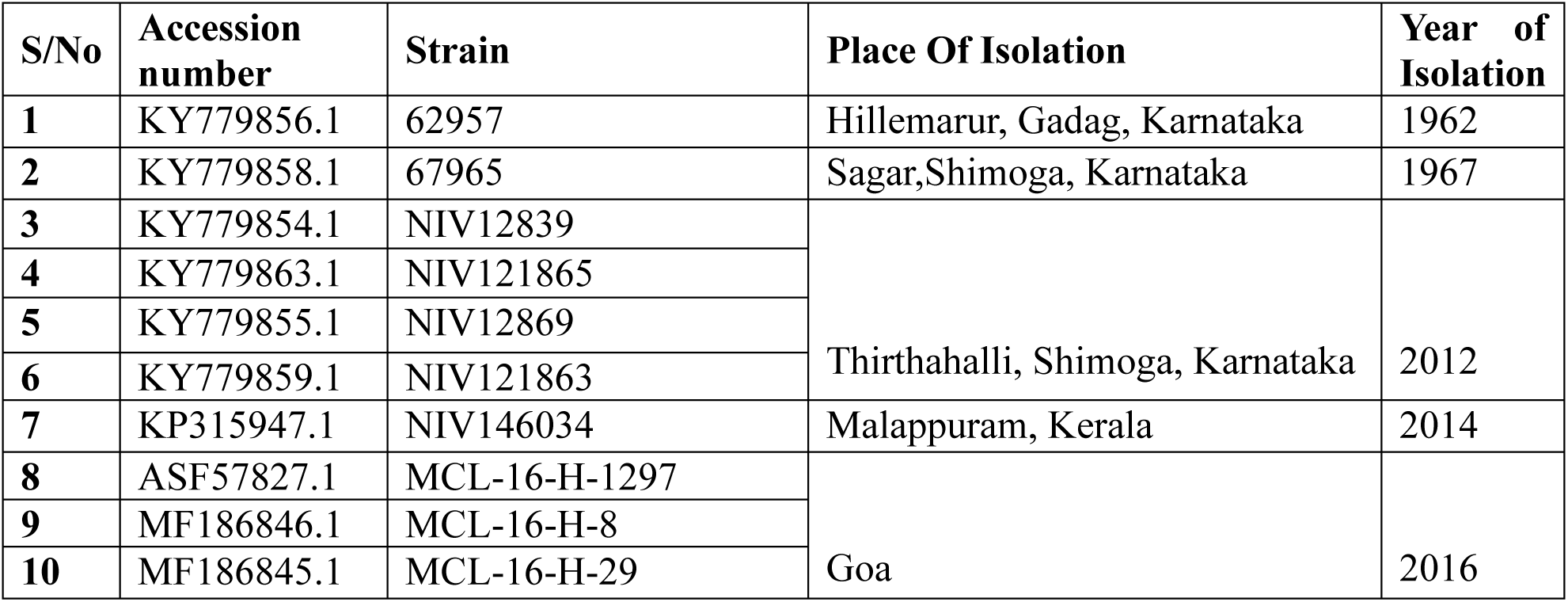
Isolates of KFD Virus shortlisted for the study.

**Table 2:**
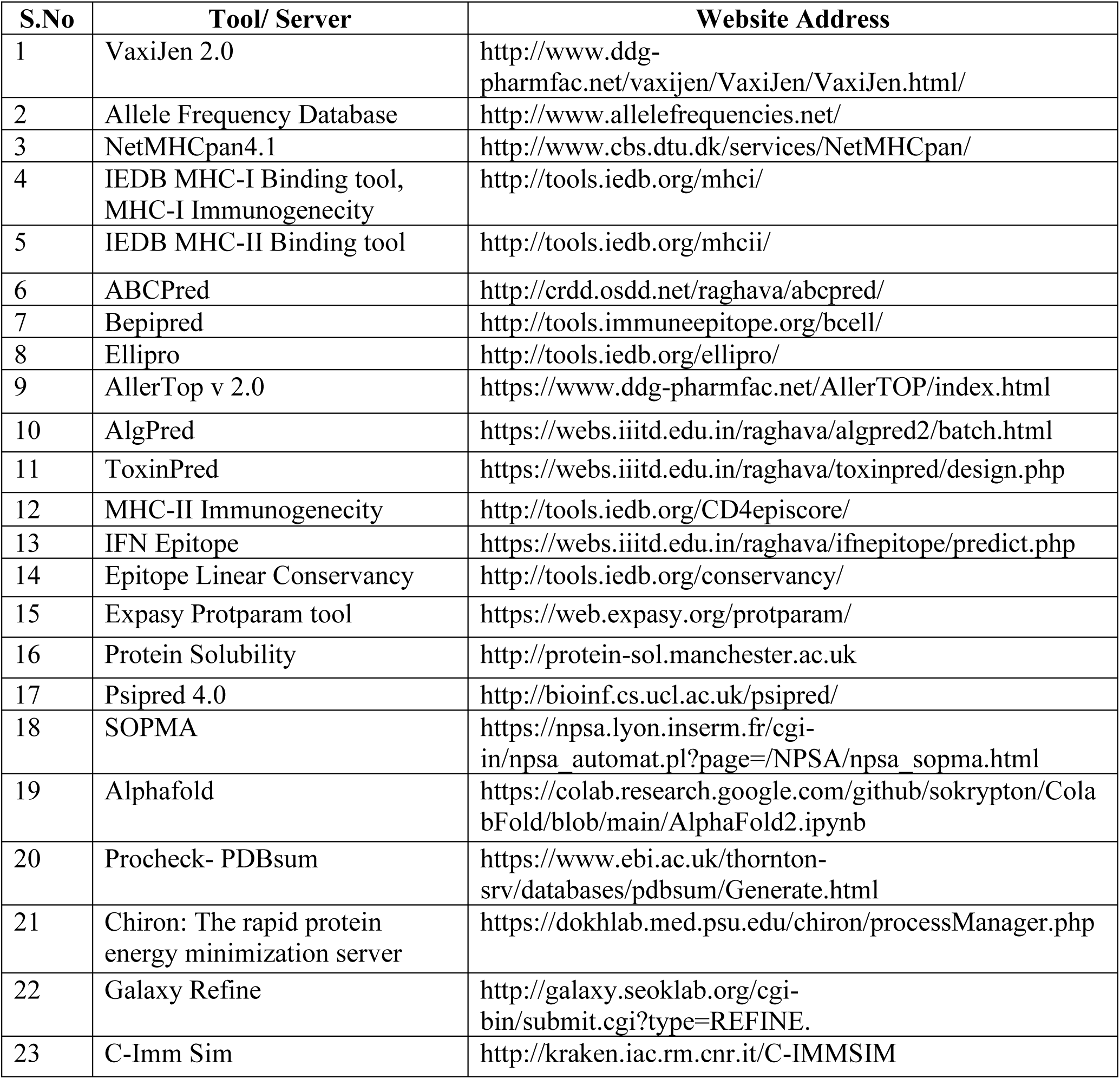
*In silico* tools or servers used in this computational research work.

### Selection of Suitable Antigenic Protein

Antigenicity pertains to the ability of a foreign entity or an antigen to be identified by specific antibodies that are produced when a substance is immune-mediated. The KFD viral protein antigenicity was determined by the VaxiJen 2.0 server at a threshold of 0.4[29]. For the epitope prediction, the FASTA of the protein sequence with the greatest antigenicity score will be selected and saved.

### Cytotoxic and Helper T Lymphocytes Epitopes’ Prediction

In order to kindle an immune response, antibodies / T-cell receptors connect with epitope. Adaptive immune responses are elicited by the T-lymphocytes. In eliciting cytotoxic-T cell (CTL) responses, antigens are processed intracellularly and linear epitopes are primarily targeted. Human Leukocyte Antigen (HLA) plays a significant role and these antigens are in charge of spotting cells as they enter the body. In order to develop a vaccine, its epitopes must attach to more than one MHC allele and cover almost all of the major populations in the world. The Allele Frequency Database was used to choose MHC Class 1 HLA-A * 24:02 and MHC Class 2 HLA-DRB1 * 15:01 due to the recurring incidence of HLA predominance in the Indian population [30], [31]. The default parameters of epitope prediction programs such as IEDB MHC-I tool and NetMHCpan4.1 were used to predict CTL epitopes of nine mers each, whereas IEDB MHC-II tool was utilised to predict HTL. 15-mer epitopes with a percentile rank less than 10 were anticipated [32]. The epitopes with the minimum percentile rank exhibit a high level of attraction of MHC-II.

### Linear B-cell Epitope Prediction

B-cell epitopes are necessary for eliciting an antibody-mediated natural defenses, which in turn kindles B cells to produce appropriate antibodies. It is predicted using tools like ABC pred, Bepipred and Ellipro from IEDB Analysis Resource [33], [34], [35]. A higher peptide score indicates a greater likelihood of becoming an epitope [36]. Residues that achieve scores above the default threshold of 0.35 are considered as an epitope[35].

### Antigenicity Prediction of Epitopes

The epitopes’antigenic nature is determined using VaxiJen2.0 at a cutoff value of 0.4[29]. This tool was employed to determine the antigenic nature of the predicted 25 CTL&HTL, and 66 B-cell epitopes.

### Allergenicity Prediction of Protein Sequences

Prior to developing a vaccination candidate, allergens must be identified. If a protein and recognised allergens share more than 35% of the same sequences over an 80 amino acid window, it is considered to be a possible cause for allergy[37]. Using AllerTop v.2.0, the anticipated epitopes’ allergic nature was identified[38]. The chosen epitopes were subjected for analysis using Allertop and the epitopes that would be found to be non-allergenic will be further used for the vaccine construct and studies. The same results were cross-checked using AlgPred [39].

### Toxicity Evaluation of Predicted T-Cell Epitopes

Toxicity test is to determine whether a substance has detrimental impacts on human health, animal health, or the environment in general. The toxic nature of the selected epitopes was predicted using the server ToxinPred [40]. The tool generates all the possible mutants of a given peptide and also identifies the toxic regions in proteins[40].

### Immunogenicity Prediction of Predicted T-Cell Epitopes

Immune cells with the capacity to develop pathogen specific memory which confer immunological defence are T and B cells, which facilitate adaptive immunity. In order for B and T-cells to become memory cells, they must recognize specific targets (antigens) on pathogens through specialized receptors. Henceforth to design effective vaccines and to have a deeper comprehension of the immune system, it is significant to precisely predict immunogenic CTL&HTL epitopes. Immunogenicity predicting tools available from IEDB were utilized to analyze the immunogenicity of the chosen epitopes. An increased likelihood of eliciting an immunological response is indicated by a higher IEDB Class-I MHC prediction score. To predict the immunogenicity score of MHC-II/CD4 epitopes, the default parameters are set with a maximum combined score threshold value of 90[41].

### Prediction of IFN Gamma-Inducing Epitopes

The IFN epitope, was used to further investigate the screened 10 HTL-epitopes for their capacity to elicit an IFN-γ immune response [42]. The IFN epitope enables users to determine the peptides or antigens that induce MHC class II binding when exposed to IFN-γ. For the *in silico* vaccine candidate development, the epitopes that showed positive IFN response outcomes were ultimately selected.

### Conservation Analysis of Predicted T-Cell Epitopes

Measures of identity and conservancy are used to define and characterize epitope or protein variability. It is important to perform a conservancy analysis when estimating epitopes, as it determines whether the epitope is cross reactive among different isolates of a virus, or with different microorganisms which have varying degrees of pathogenicity. We computed epitope linear conservancy across antigens using the Epitope Analysis Tools of IEDB which computes the conservation degree of an epitope within a range of protein sequences at a particular identity level. The tool was employed to shortlist the epitopes with a criterion that is higher than or equal to 100% sequence identity [43].

### Conceptualization of Multi-Epitope PKFDVac-I (PIIC Vac Candidate-I)

The MEPV sequence was meticulously formulated utilizing innocuous, non-allergenic, and significantly antigenic Cytotoxic T lymphocytes, B-cell epitopes, and Helper T lymphocyte epitopes. These epitopes were interconnected via GPGPG, AAY, and KK linkers to guarantee that the vaccine construct functions as a distinct immunogen and invokes greater concentrations of antibody production than a solitary immunogen [44].By exploiting AAY as a proteosome cleavage site, protein’s stability, immunogenicity, and epitope presentation can be altered[45]. KK is used to maintain the immunogenicity of the vaccine constructs[46]. Figure 1 provides a schematic representation of the complete process involved in predicting epitopes and developing PKFDVac-I.

**Figure 1:**
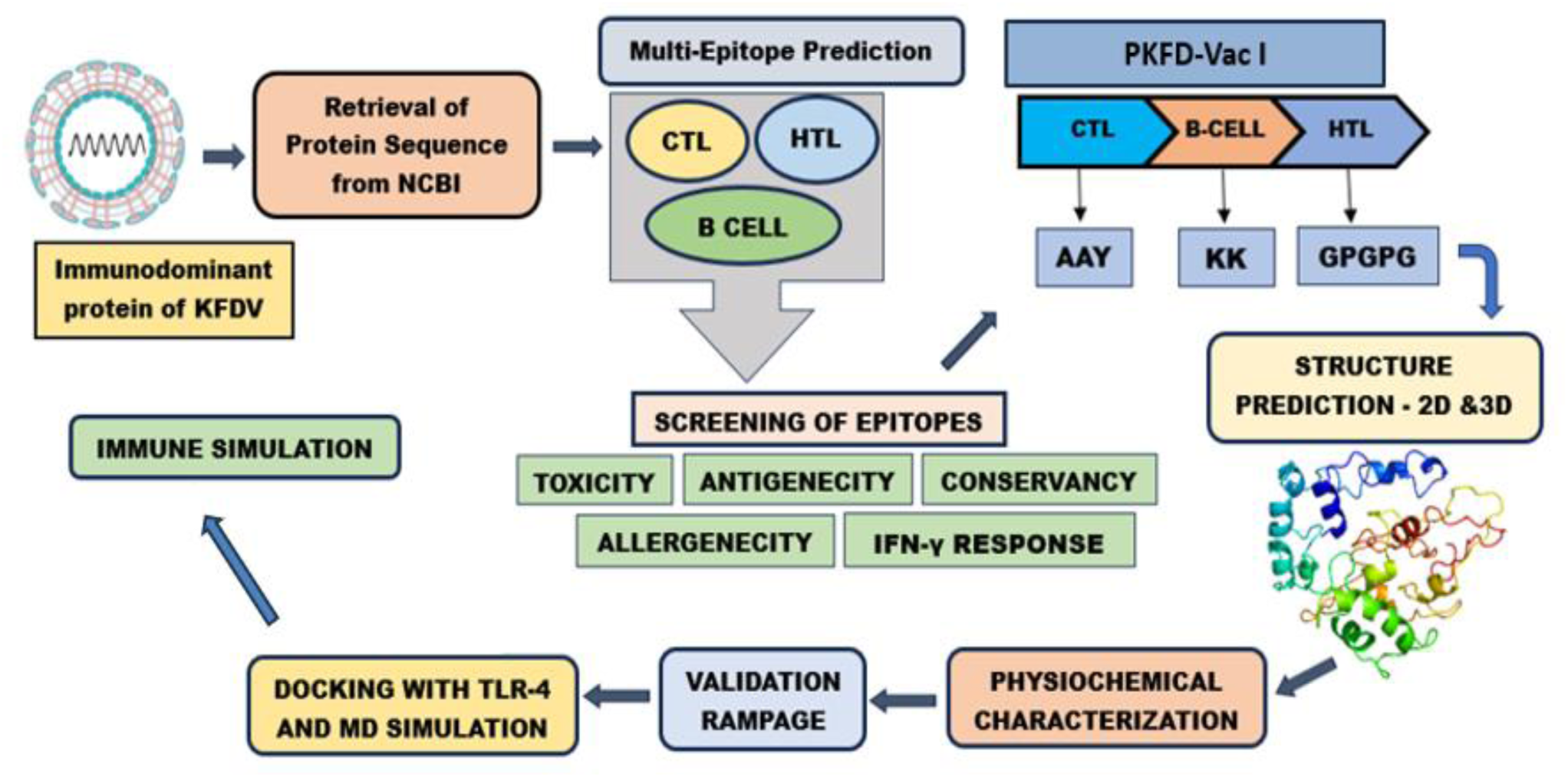
Comprehensive Workflow for Epitope Prediction and Development of PKFDVac-I.

### Physiochemical Properties of PKFDVac-I

Vaccine quality attributes of PKFDVac-I were analyzed along with other physiochemical characteristics. The antigenicity was determined using VaxijJen v2.0 at a threshold value of 0.4. The allergenic property was determined using AllerTopv.20[38]. Employing the Expasy Protparam tool, the physiochemical attributes were forecasted [47]. The solubility of the MEPV was evaluated utilizing the Protein-Sol server [48].

## 2D and 3D Structure Prediction

The 2D structure of PKFDVac-I was determined using PSIPRED and SOPMA servers. The online tool Psipred 4.0 accurately predicts the transmembrane helix, fold, domain recognition, and topology [49]. SOPMA server uses information from a multiple sequence alignment (MSA) of a protein belonging to the same family to predict the 2D structure [50].

3D structure of PKFDVac-I was modeled using Alphafold colab It is the first computational method that routinely predict protein structures with atomic exactness even when there is no known structure identical to it [51].

### Refinement of the 3Dstructure and Validation

Validation of 3Dstructure is a crucial phase as it identifies potential boo-boos in models that were predicted. A software PROCHECK with a visual database called PDBsum [52] gives a quick overview of the information contained in each 3D structure that has been submitted in PDB. The PROCHECK programmes are helpful for evaluating the excellence of protein structures, both those that have already been solved and those that are being modelled after existing structures.

The model which showed a better score in the Ramachandran plot statistical analysis will be refined and energy minimized. The minimization of the energy of the 3D model was done using Chiron: The rapid protein energy minimization server [53] and upgraded using the Galaxy Refine web server.

### Molecular level Docking of the PKFDVac-I Vaccine Candidate Construct with the TLR-4 Immune Receptor

The concept of Molecular Docking is based on the understanding that an effective immune response depends on the interaction between an antigen and a specific immune receptor. To examine the interactions between vaccines and receptors, researchers utilized the Maestro program from the Schrödinger suite to conduct a molecular docking study [54]. The tertiary structure of PKFDVac-I was used as a ligand, and the receptor TLR4 (PDB: 4G8A) was downloaded from PDB. The vaccine model and receptor were prepared with Schrödinger Protein Preparation Wizard using default settings. During the process, the hydrogen atoms were added, crystals were removed, and fractional charges were allocated using the OPLS-2005 force-field. The binding site of Chain C (MD2) was found to be an active binding site. A grid box measuring (96 x 96 x 96 Å) was selected to surround the Chain-C area, with the grid center located at the center of this previously described binding site. The docking was performed with a Protein-Protein docking module. The top-docked poses were selected as the lowest Glide score. The PDBsum online server was utilized for the analysis of the binding residues and interaction surfaces of the TLR4 and vaccine complexes.

### Molecular Dynamics Simulation

To further examine the interaction between vaccine-receptor complexes, a Molecular Dynamics (MD) simulation was conducted. The simulation was executed using Schrödinger’s Desmond module v2.3. The System Builder application within Desmond was utilized to prepare the systems for subsequent calculations. An orthorhombic periodic boundary box with a 10 Å buffer distance on all sides of the reservoir was designated to ensure a specific volume. Following the solvation of the protein-vaccine complex within this system, energy minimization and relaxation were carried out using the default protocol available in the Desmond module, employing the OPLS 2005 force field parameters [55]. The simulation was run for 100ns at 300K in an NPT ensemble, with total energy (kcal/mol) recorded every 100 ps. Upon completion of the MD simulation, the stability of the complex was evaluated using Desmond’s Simulation Event Analysis tool. This tool analyzed the trajectory file produced by the MD simulation to determine hydrogen bond interactions, RMSF, and RMSD.

### Immune Simulation

To assess the immunological retort of the engineered peptide, immune simulation using the C-Imm sim server was carried out [56]. In pre-clinical testing of the prototype vaccine, shorter intervals between doses are used to quickly elevate antibody titers above protective levels. Clinically, the recommended minimum period between two vaccine doses is one month, or approximately four weeks[57]. For our simulation, we used time step parameters of 1, 84, and 168, corresponding to 8-hour intervals over 1000 simulation steps[58][59] The immune response simulation followed the same protocol and default parameters as reported in previous studies. The capability of the PKFDVac-I multiepitope peptide vaccine to stimulate immune system cells, including dendritic cells, immunoglobulins, B-cell lymphocytes, HTL, CTL, and natural killer cells, was studied.

## RESULTS

### Identification, Analysis of KFD Viral Strain and Retrieval of Immunodominant Protein

The chosen Indian strains’ accession codes and the gene sequence encoding their envelope protein were obtained from NCBI and saved in FASTA format. The glycoprotein E of KFDV consisted of 496 aminoacids. To date, all tick-borne Flaviviruses analyzed have three potential N-glycosylation sites in the E protein[60]. It helps create a bridge between the host and viral cellular membranes, allowing the virus entry to the host cell. In addition, it induces host immunity by producing neutralizing and protective antibodies[61]. Moreover, structural components of the E protein aid in viral attachment, fusion, hemagglutination, host range, penetration, and cell tropism, as well as viral virulence and attenuation during spontaneous infection or vaccination [62]. E protein plays a significant role in determining virus virulence or attenuation in studies with neutralization-resistant escape mutants and attenuated strains of different Flaviviruses [63] and vaccine development[23][26].

### Antigenic evaluation of Immunodominant protein of KFDV

The antigenicity score was analyzed and a comparative study (Table 3) found that all the strains gave an antigenicity score of above 0.45 and the strain with the highest antigenic value was selected for the further study. Out of the protein sequence of the ten isolates, strain 62957 and 67965 isolated from humans showed the highest antigenicity score of 0.6649, but to have a species isolated and sequenced recently, MCL-16-H-1297 (0.6634) isolated in the year 2016 was selected than the ones isolated from 1962 and 1967 respectively.

**Table 3:**
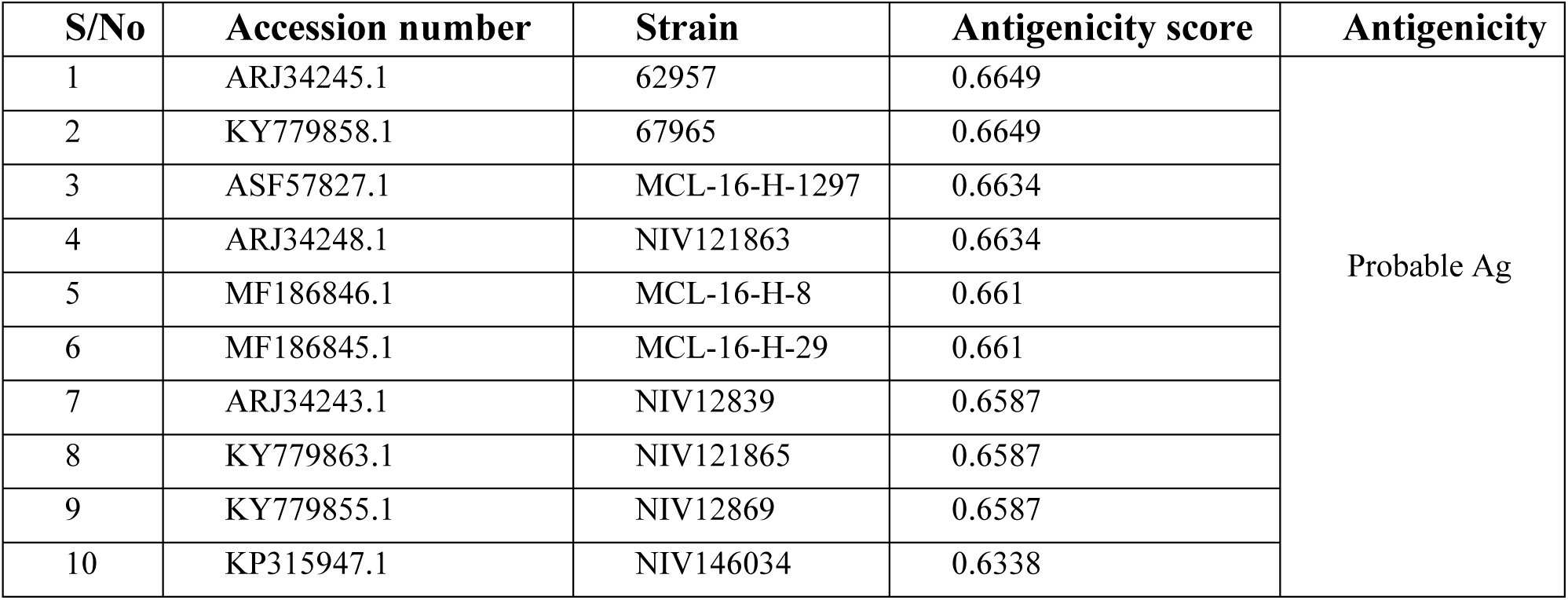
Estimated antigenicity of the chosen isolates’ E-protein.

| S/No | Accession number | Strain | Antigenicity score | Antigenicity |
| --- | --- | --- | --- | --- |
| 1 | ARJ34245.1 | 62957 | 0.6649 | Probable Ag |
| 2 | KY779858.1 | 67965 | 0.6649 |  |
| 3 | ASF57827.1 | MCL-16-H-1297 | 0.6634 |  |
| 4 | ARJ34248.1 | NIV121863 | 0.6634 |  |
| 5 | MF186846.1 | MCL-16-H-8 | 0.661 |  |
| 6 | MF186845.1 | MCL-16-H-29 | 0.661 |  |
| 7 | ARJ34243.1 | NIV12839 | 0.6587 |  |
| 8 | KY779863.1 | NIV121865 | 0.6587 |  |
| 9 | KY779855.1 | NIV12869 | 0.6587 |  |
| 10 | KP315947.1 | NIV146034 | 0.6338 |  |

### Cytotoxic and Helper T Lymphocytes Epitopes’ Prediction

Using *in silico* tools, the epitopes were predicted and selected based on the percentage rank. A higher score denotes a higher likelihood of triggering an immunological response and the top 25 CTL epitopes were carefully chosen for further study (Table 4). The lower adjusted rank of the epitopes in IEDB indicates higher binding of epitopes. Based on the MHC-II (HTL) binding percentile rank, top 25 epitopes were selected (Table 5).

**Table 4:**
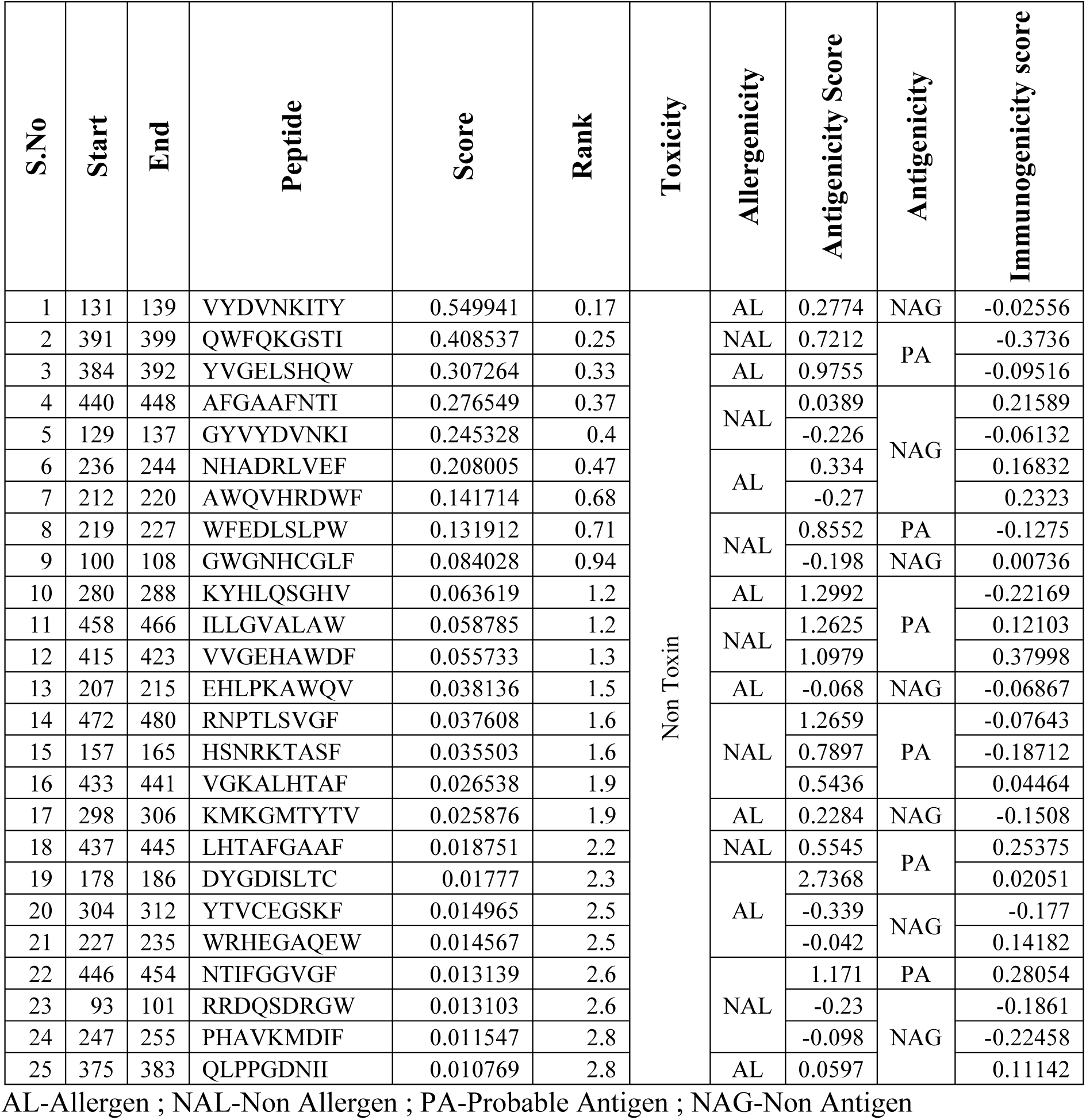
MHC-I Binding prediction results and analysis report highlighting the Toxicity, Allergenicity, Antigenicity and Immunogenicity status of the predicted CTL epitopes.

**Table 5:**
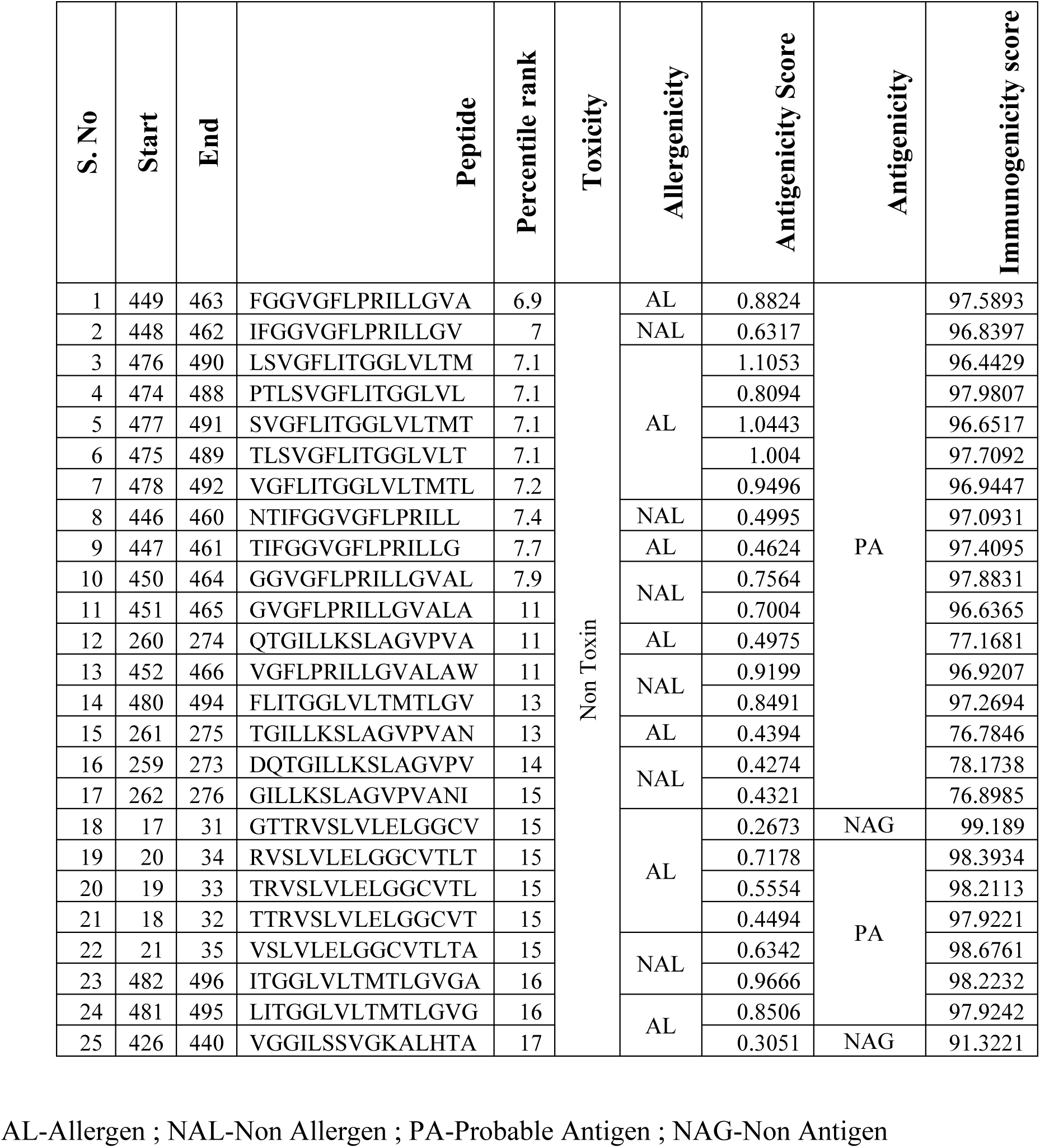
MHC-II Binding prediction results and analysis report highlighting the Toxicity, Allergenicity, Antigenicity and Immunogenicity status of the predicted HTL epitopes.

### Linear B-cell Epitope Prediction

The 496 amino acid long KFDV E protein sequence when submitted in the ABCPred server with a threshold setting of 0.51, a total of 52 B-cell epitope (S.No 1-52) of 16 mer were predicted, 10 epitopes of 16 mer (S.No 53-63 ) were predicted using the Ellipro tool and 3 epitopes of different lengths were predicted using the IEDB Bepipred tool (S.No 64-66). Since a peptide with a higher score has a greater chance of being chosen as an epitope, the 66 epitopes (Table 6) were arranged in descending order. In order to develop the vaccine candidate, the epitope with the highest score was selected[36].

**Table 6:**
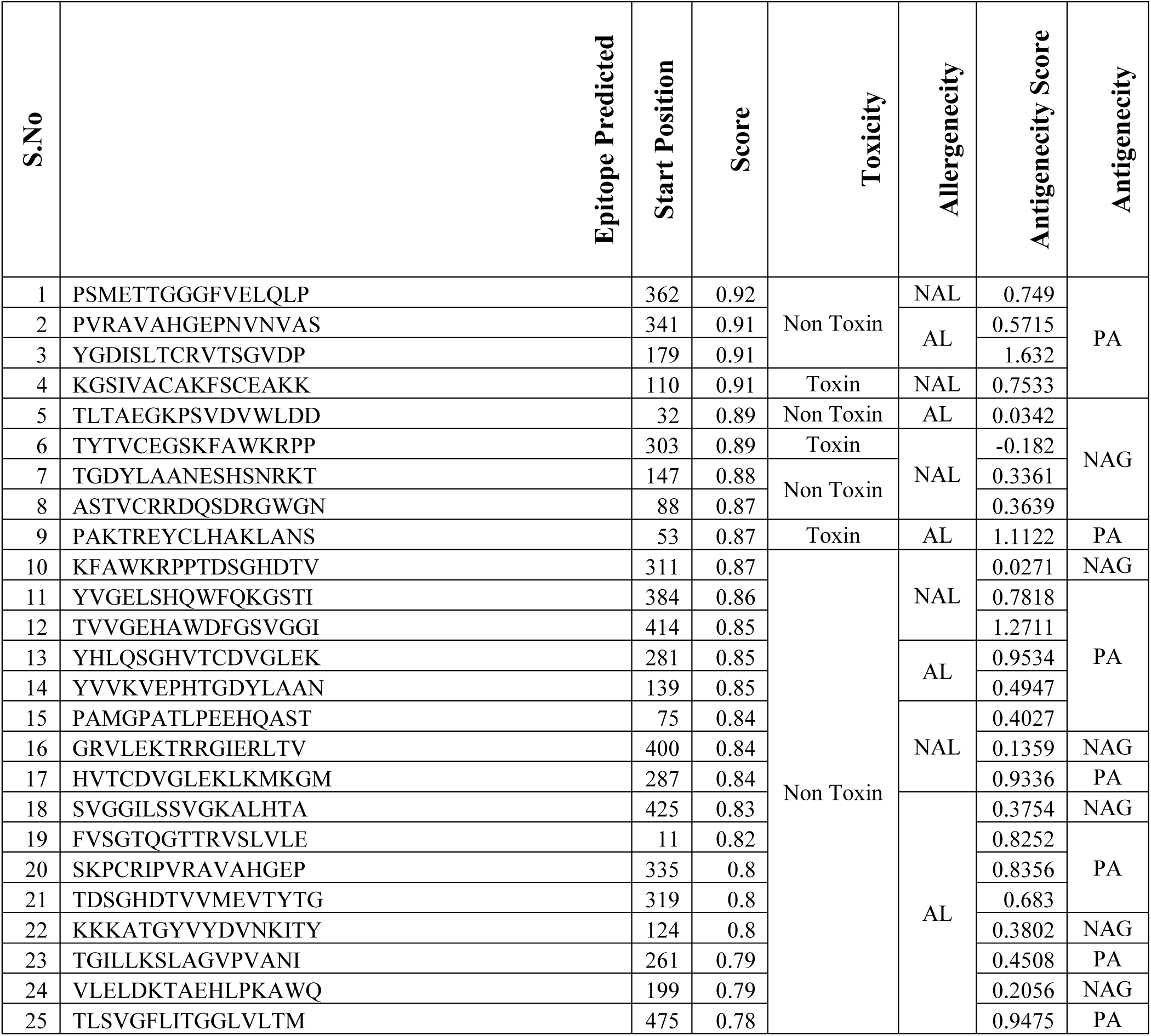
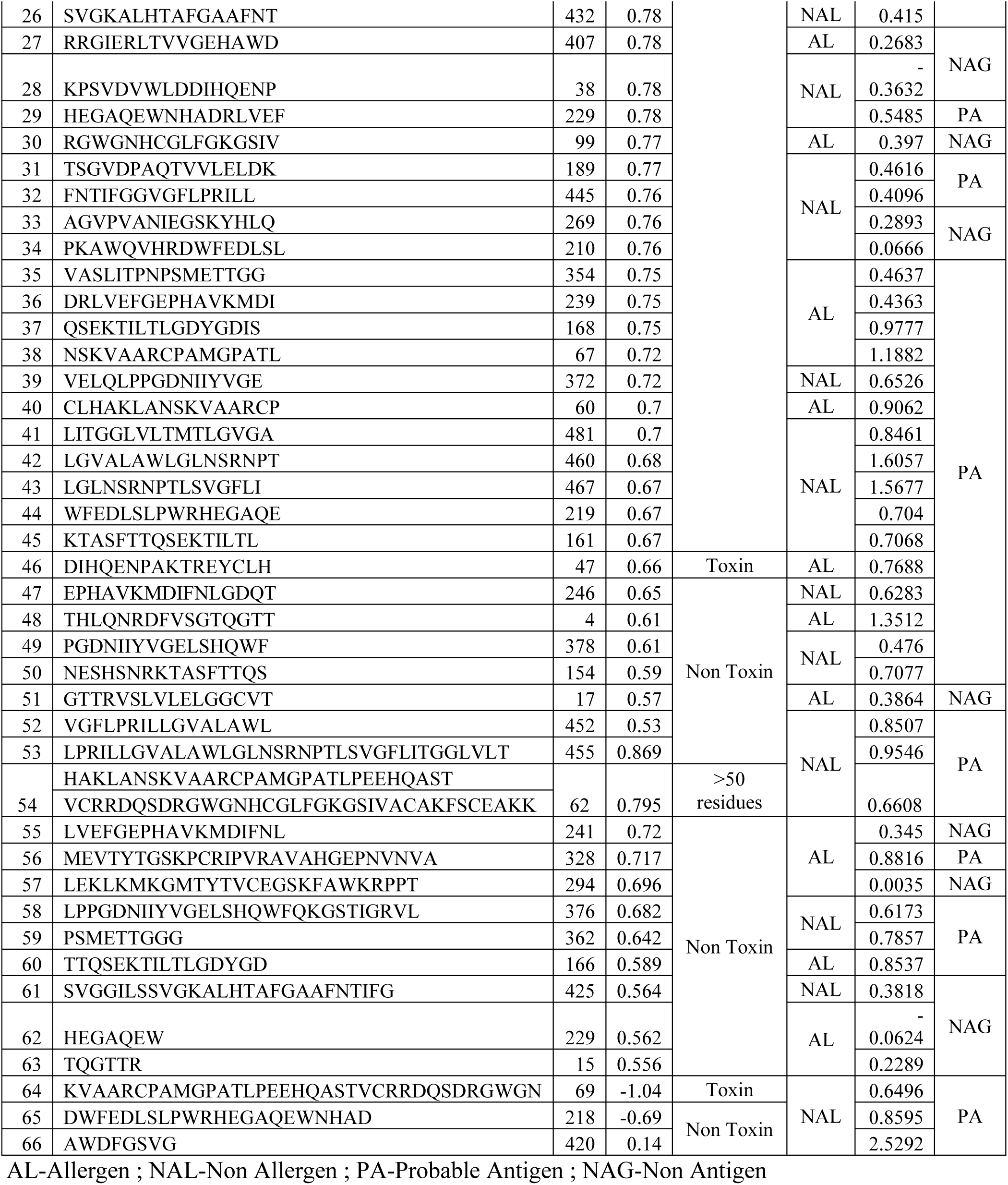
B-cell Binding prediction results and report of the Toxicity, Allergenicity and Antigenicity analysis.

### Epitopes’ Antigenicity Prediction

The antigenic nature of the predicted epitopes at a threshold value of 0.4 was determined and compared. It was found that out of the 25 epitopes predicted, only 12 CTL & 23 HTL epitopes were found to be antigenic (Table 4 and 5). Out of the 66 B-cell epitopes predicted, only 46 can be considered as probable antigen (Table 6).

### Epitopes’ Allergenicity Prediction

For the 25 CTL and HTL epitopes, allergenicity predictions were made. According to Tables 4 and 5, respectively, 10 HTL and 14 CTL epitopes were non-allergenic. 36 of the 66 B-cell epitopes turned out to be non-allergic (Table 6).

### Epitopes’ Immunogenicity Analysis

The high-binding-ability epitopes that were chosen are subsequently put through additional testing to forecast their immunogenicity. Positive epitope scores are indicative of high immunogenicity, which denotes that they have a strong capacity to activate T cells and produce cellular immunity. In Tables 4 and 5, the predicted immunogenicity scores are displayed.

### Epitopes’ Toxicity Analysis

The selected epitopes were subjected for toxicity prediction at a threshold set at 0.5. The results obtained showed that none among the chosen CTL and HTL epitopes exhibited toxicity (Table 4 and 5), but 5 B-cell epitopes were toxic in nature (Table 6).

A vaccine candidate must be highly immunogenic, highly antigenic, non-toxic, and non-allergic in order to be effective. Only nine CTL epitopes and ten HTL epitopes are compatible with the aforementioned, according to the data in Tables 4 and 5. Only 24 B-cells (Table 7) are shortlisted for the vaccine candidate build and additional research because they fit that description, as Table 6 demonstrates.

**Table 7:** Selection of final B-cell epitopes for vaccine construct based on antigenicity scores and prediction tools.

| S. No | Epitope Predicted | Antigenicity Score |
| --- | --- | --- |
| 1 | LGVALAWLGLNSRNPT | 1.6057 |
| 2 | LGLNSRNPTLSVGFLI | 1.5677 |
| 3 | TVVGEHAWDFGSVGGI | 1.2711 |
| 4 | HVTCDVGLEKLKMKGM | 0.9336 |
| 5 | VGFLPRILLGVALAWL | 0.8507 |
| 6 | LITGGLVLTMTLGVGA | 0.8461 |
| 7 | YVGELSHQWFQKGSTI | 0.7818 |
| 8 | PSMETTGGGFVELQLP | 0.749 |
| 9 | NESHSNRKTASFTTQS | 0.7077 |
| 10 | KTASFTTQSEKTILTL | 0.7068 |
| 11 | WFEDLSLPWRHEGAQE | 0.704 |
| 12 | VELQLPPGDNIIYVGE | 0.6526 |
| 13 | EPHAVKMDIFNLGDQT | 0.6283 |
| 14 | HEGAQEWNHADRLVEF | 0.5485 |
| 15 | PGDNIIYVGELSHQWF | 0.476 |
| 16 | TSGVDPAQTVVLELDK | 0.4616 |
| 17 | SVGKALHTAFGAAFNT | 0.415 |
| 18 | FNTIFGGVGFLPRILL | 0.4096 |
| 19 | PAMGPATLPEEHQAST | 0.4027 |
| 20 | LPRILLGVALAWLGLNSRNPTLSVGFLITGGLVLT | 0.9546 |
| 21 | PSMETTGGG | 0.7857 |
| 22 | LPPGDNIIYVGELSHQWFQKGSTIGRVL | 0.6173 |
| 23 | DWFEDLSLPWRHEGAQEWNHAD | 0.8595 |
| 24 | AWDFGSVG | 2.5292 |

### Prediction of IFN-γ Inducing Epitopes

Out of the 10 shortlisted HTL epitopes, only 3 were found to be positive for IFN-**γ** (Table 8), and hence chosen for the vaccine construct.

**Table 8:**
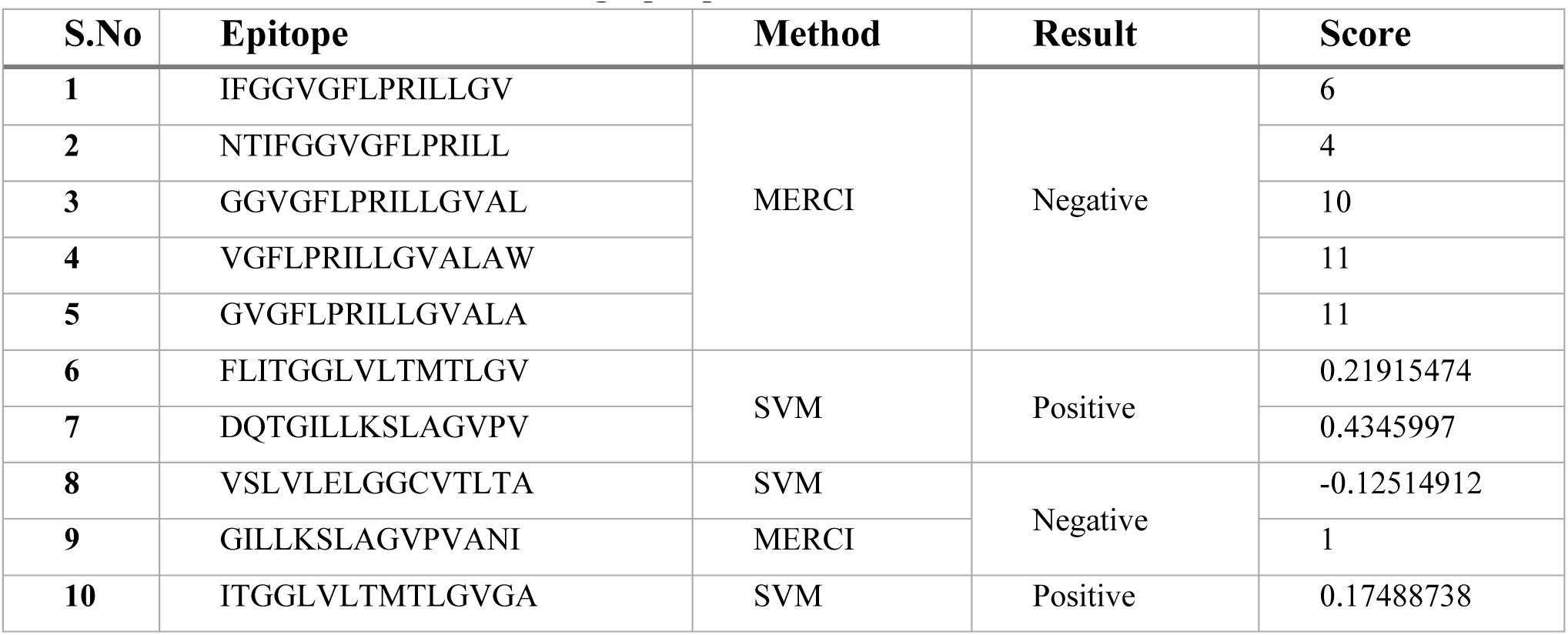
Gamma Interferon Inducing Epitopes.

### Conservancy Analysis

Conservancy analysis of the carefully chosen T-cell epitopes predicted from the Immunodominant protein E of the KFD virus was carried out and it is necessary to align a protein epitope with every protein sequence within a given set of sequences to determine how conserved that epitope is. To achieve effective immunization, epitopes must be conserved among strains all over the world. The predicted result shows 100% conservancy for all the selected epitopes isolated from the strain MCL-16-H-1297 across the other shortlisted strains of the KFD virus used in the study.

### Construction of the Multiepitope Vaccine Candidate PKFDVac-I Using the Selected Epitopes

5CTL, 3HTL, and 8 B-cell epitopes were finalized from the predicted and screened candidates. These were then linked together to form the vaccine construct, PKFDVac-I, with a total length of 279 amino acids. Figure 2 illustrates the schematic representation of this vaccine construct.

**Figure 2:**
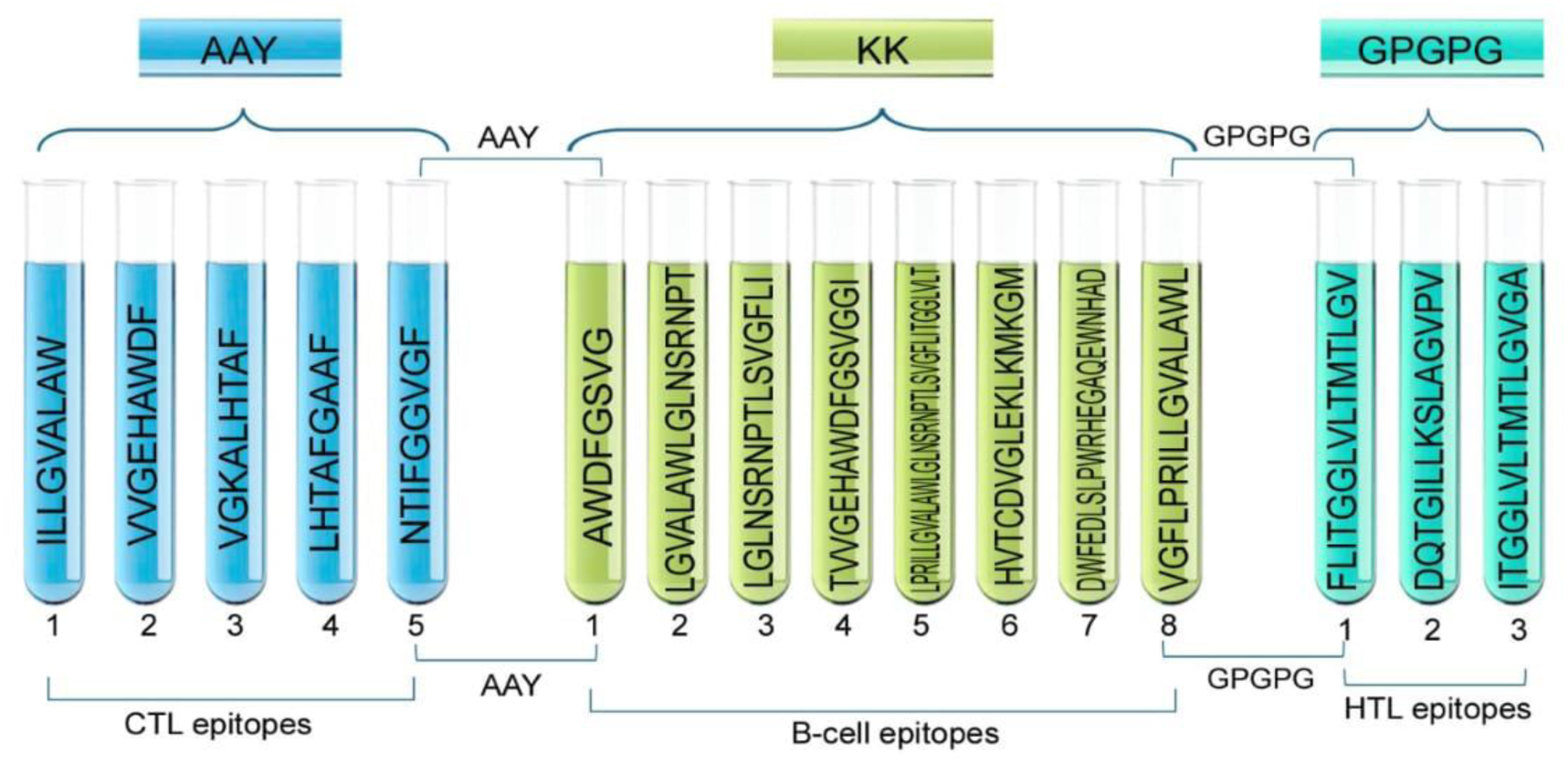
**Schematic Representation of PKFDVac-I, the Multiepitope Peptide Vaccine Candidate.**

### Physiochemical Properties and Solubility Prediction of PKFDVac-I

The protective antigenic score and allergenicity of PKFDVac-I was 0.8882 and −0.81924 respectively, which reveals it as a probable antigen which is non allergenic based on their amino acid composition [38]. The physiochemical properties i.e., molecular weight of PKFDVac-I comprising of 279 amino acid was found to be 29162.32 Daltons. Proteins under 110 kDa are considered better as they can be purified and used for vaccine development more easily [64].

The pI was found to be 9.79. The bulk of the amino acids in the peptide structure have basic structures, evidenced by the comparatively high pI[65]. The atomic composition was elucidated with the chemical formula C1363H2128N350O349S5 comprising of 4195 atoms. Assuming that all pairings of Cys residues form cystines, the Ext. coefficient was calculated as 62450 Abs 0.1% (=1 g/l) 2.141. The half-life was found to be 20 hours for human reticulocytes in lab conditions, 30 minutes for yeast in living conditions, and over 10 hours for *Escherichia coli* in living conditions. The Instability Index (II) is 11.80, which means the protein is stable, with an Aliphatic index of 108.39 and a GRAVY of 0.448. A high aliphatic index indicates good thermal stability, as demonstrated by the new vaccine. The protein appears to be hydrophilic and soluble based on the low GRAVY[65]. Since some amino acids, like alanine, valine, cysteine, and methionine, were found to be under-represented in antibody-antigen recognition sites while aromatic residues were found to be over-represented, the amino acid composition of PKFDVac-I was extracted to ascertain the percentage of aliphatic and aromatic amino acids present in it [66]. The quality and physiochemical attributes of PKFDVac-I is summarized in Table 9.

**Table 9:**
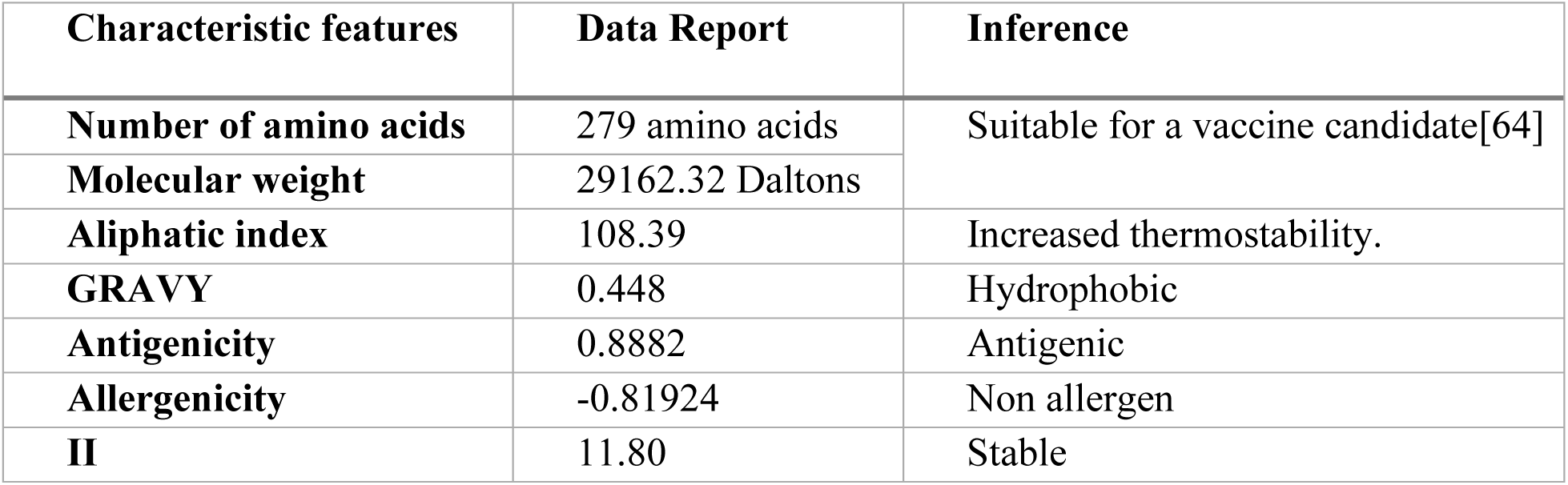
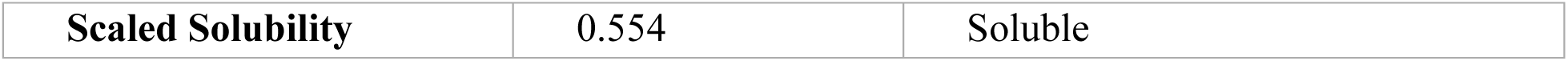
Quality attributes of PKFDVac-I.

The data obtained from Expasy Protparam showed that the aliphatic amino acid contributes to 58.9% and the aromatic to 10.1% of the overall amino acid composition. The PsiPred[67] data obtained while predicting the 2D structure showed that the PKFDVac-I comprised of small non polar (41.21 %), hydrophobic (28.67 %), polar (20.07 %) and aromatic plus cysteine (10.03 %) group of amino acids and is represented in Figure 3.

**Figure 3:**
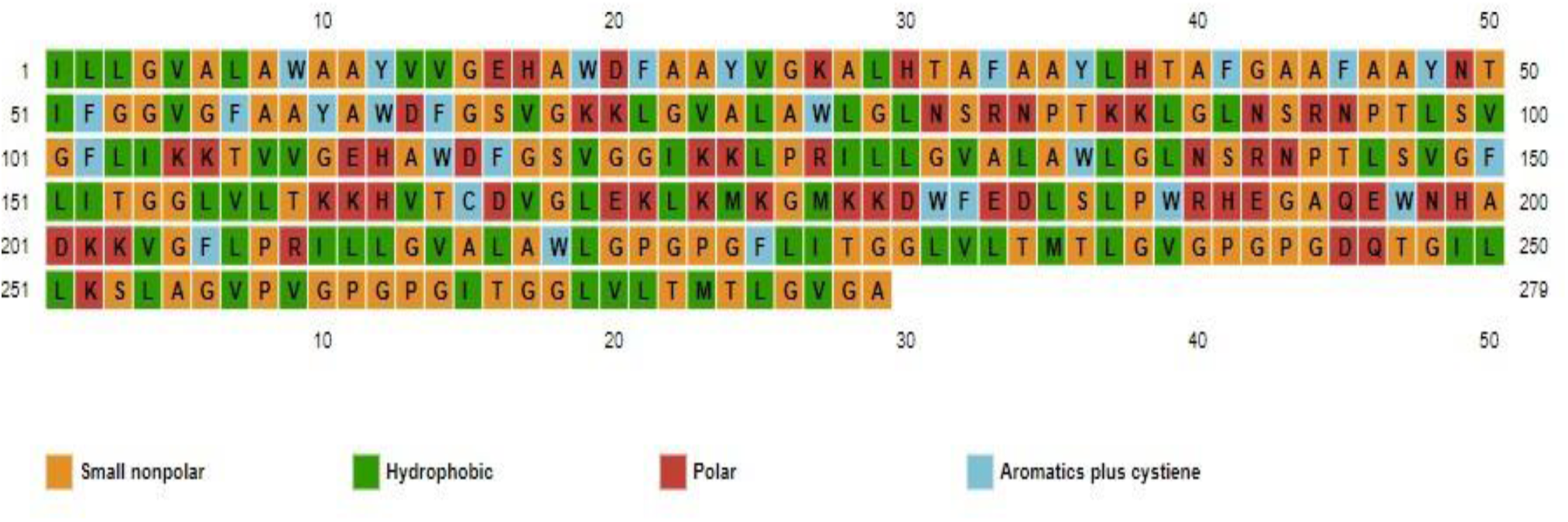
**Secondary structure predicted by PSIPRED depicting the amino acid types.**

### Secondary Structure Prediction of PKFDVac-I

The 2D structures predicted using PsiPred[67] and SOPMA is shown in Figure 4(A) and 4(B). SOPMA revealed 40.50 % coiled loops, 25.45 % β sheets and 34 % α helixes.

**Figure 4:**
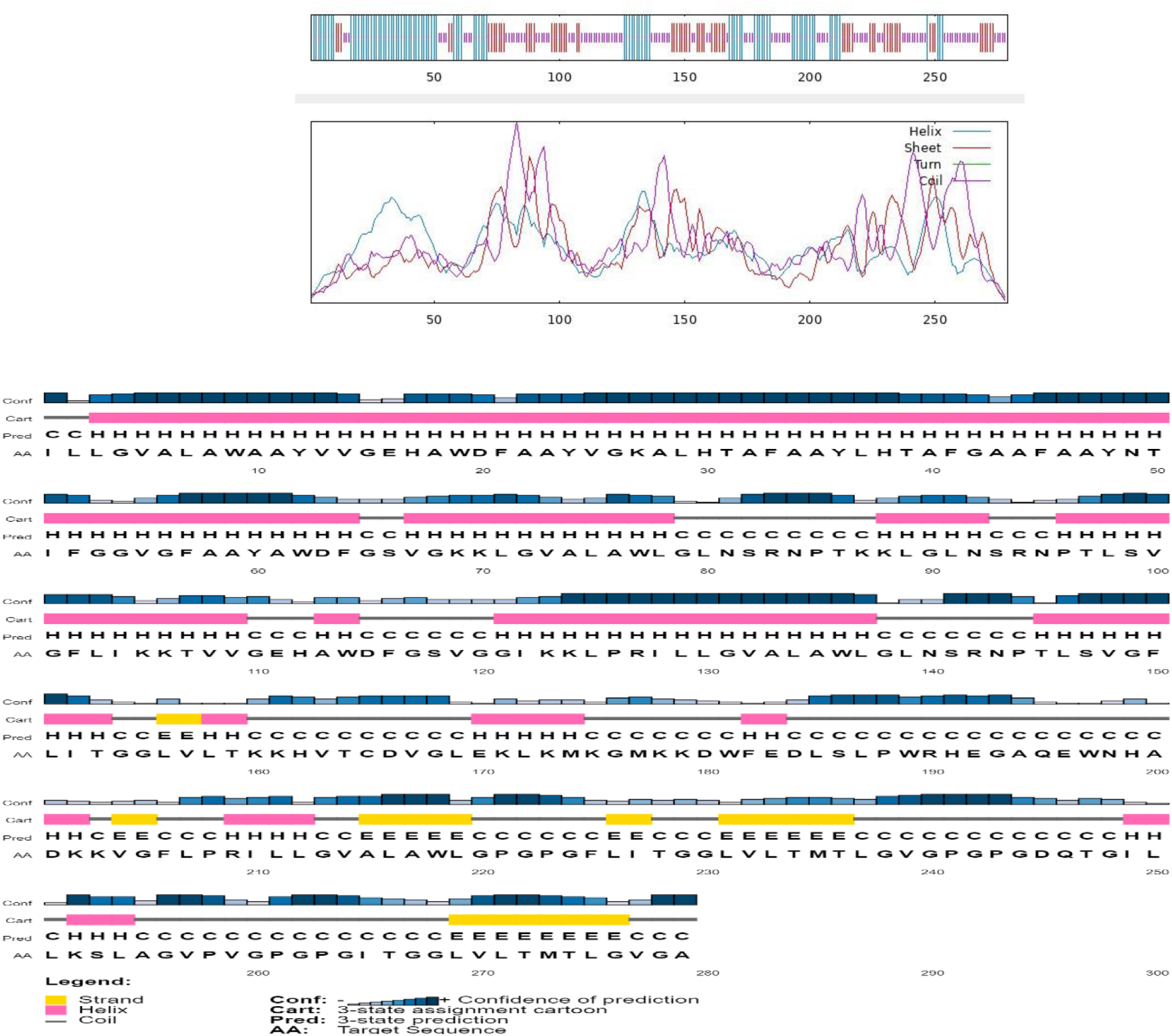
**(A) Predicted 2D structure of PKFDVac-I. (B) Graphical representation of the 2D structure of PKFDVac-I.**

### Tertiary Structure Prediction of PKFDVac-I

3D Model of PKFDVac-I was predicted with related proteins as the raw input. Five distinct 3D structural models were obtained and the greater the confidence score, better the model[51]. The structure with the highest ranking and greatest prediction score was chosen for further refining and validation[68].

### Refinement of the 3D Structure and Validation

When 3D struc-refinement was done, it was found that VDW(Van der Waals) repulsion energy was minimized to 37.3358 kcal/mol from 173.305 kcal/mol where the number of contacts between the atoms was increased to 2274 from 2150 and the number of clashes reduced to 53 from 117.The PDB structure of the refined model when further subjected for energy minimization using Galaxyrefine web server, five models were predicted with least energy score. The highest Rama favoured score, largest RMSD values and the lowest energy scores were used to determine which changed structure should be downloaded so that it can be finalized for further studies[46]. According to the results obtained, the amino-acid residues in the Rama-favored regions of the improved model Figure: 5(B)-6(B) and the original model Figure 5(A)-6(A) were 98.6% and 77.3 %, respectively. The modified structure’s Ramachandran Plot is shown in Figure 6(B), highlighting the improvements made during the refining process. The enhanced structure exhibits more stability, a prerequisite for any potential vaccine candidate, as it is critical to predictability and synthesizability[65].

**Figure 5:**
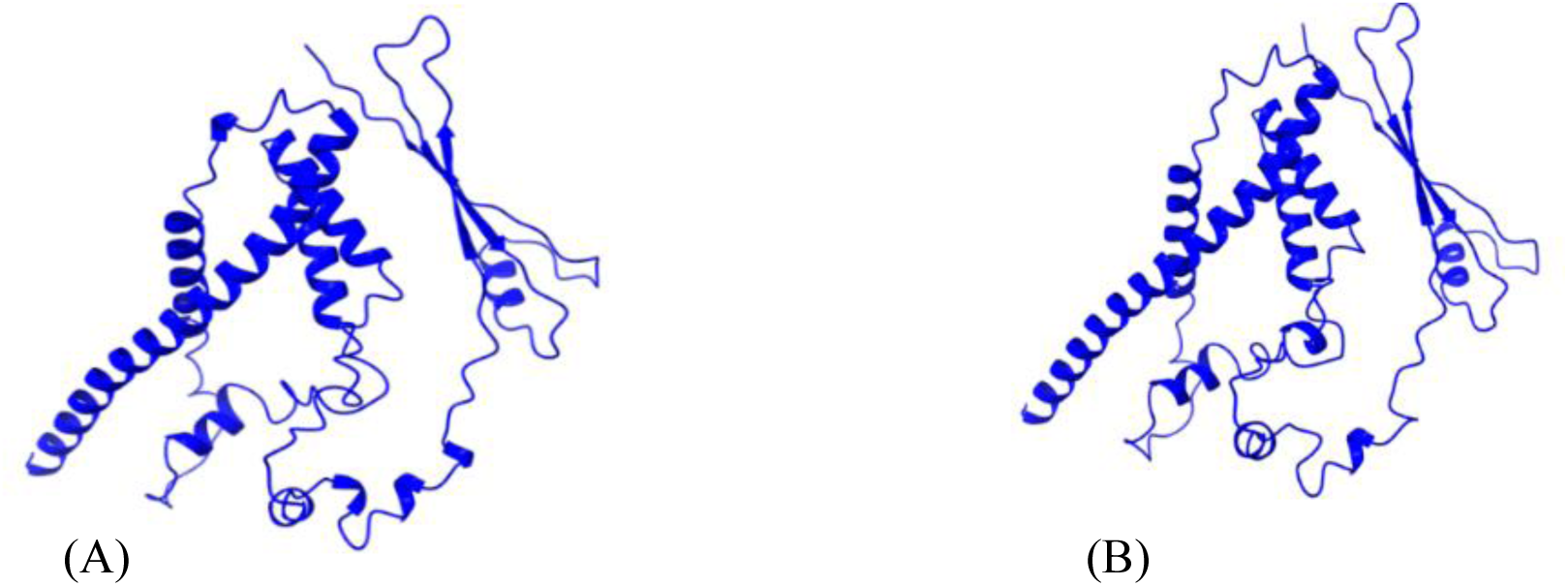
**(A) 3D structure of PKFDVac-I (B) 3D structure of PKFDVac-I obtained from Galaxy refine server after the energy minimization and refinement of the structure in Figure 5(A).**

**Figure 6:**
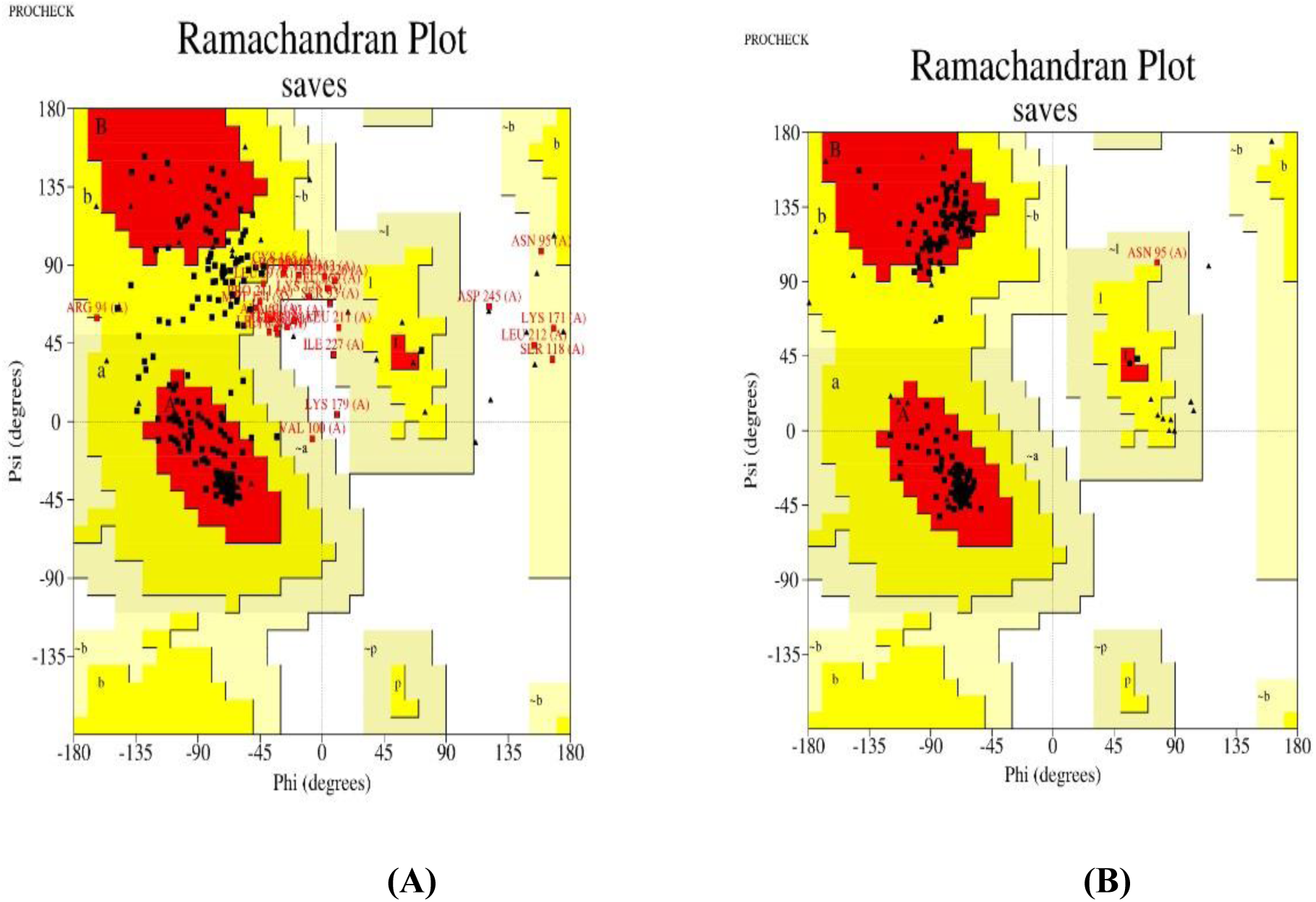
**Ramachandra plots of (A) the initial structure and (B) the refined structure of PKFDVac-I. Residues are very weakly concentrated in the favored area, as the original plot illustrates, but a greater concentration is seen in the improved plot.**

### Molecular Docking Study

Protein-protein docking was used to determine the stability of the vaccine candidate PKFDVac-I protein receptor complex. The Shrödinger server displayed 30 distinct binding mode poses. On investigating the binding affinity of the vaccine-receptor complex, the top pose with a minimum binding free energy score of **-579.716**, and Piper energy was **-1150.78** was selected. Scores are derived from potential energy changes that occur when proteins and ligands come together. Therefore, a strong binding is indicated by a very negative score, whereas a weak or nonexistent binding is indicated by a less negative or even positive score [69].The complex was then examined through binding interaction residues. The two hydrogen bonds were formed between the Lys91 and Cys95 amino acid residues with the Ala59 residue of the receptor protein and is represented in Table: 10 and Figure 7. There are no salt bridges formed between two oppositely charged residues and also no disulfide bonds formed between two cysteine residues. These interactions contribute to vaccine candidate’s stability by conformational holding of the protein. There was a total of 387 nonbonded contacts which was formed.

**Figure 7:**
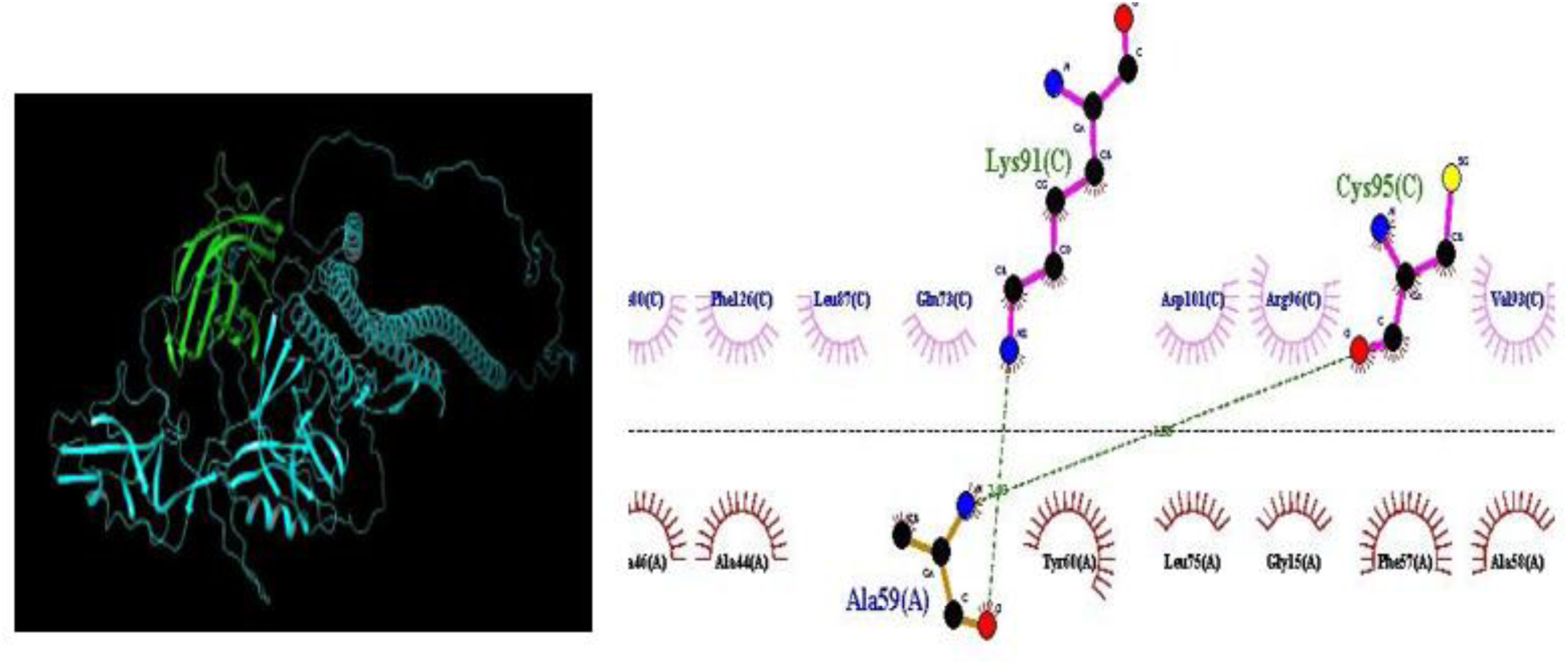
**Molecular docking interactions between PKFDVac-I and the TLR4 receptor.**

**Table 10:** Statistics of PKFDVac-I docked with TLR-4 receptor.

| S. No | Compound | Piper Score | Piper Energy | Interaction Residues |  |
| --- | --- | --- | --- | --- | --- |
|  |  |  |  | Protein | Peptide |
| 1 | PKFDVac-I TLR4 complex | -579.716 | -1150.78 | Lys91, Cys95 | Ala59 |

### Molecular Dynamics Simulation

The Desmond v2.3 package of Schrodinger software was used to run MD simulation for 100 ns. The vaccine-receptor complex’s stability was examined using RMSD and RMSF.

### RMSD calculations

RMSD is the quantitative measure of the similarity among two protein structures[70]. The graphical plot displays the developed vaccine on the right X-axis and the TLR4 protein’s RMSD on the left y-axis.

Initially, the complex was more fluctuated up to 20ns. A slight fluctuation was observed from 20ns to 40ns of the simulation time period. Furthermore, after 40ns, the simulation was stabilized, which remained consistent till 100 ns (simulation end duration). The overall RMSD plot (Figure 8) of the complex suggests that the complex initially fluctuated and stably bounded to the TLR4 binding site and remained in the constrained position until the 100 ns simulation time was over.

**Figure 8:**
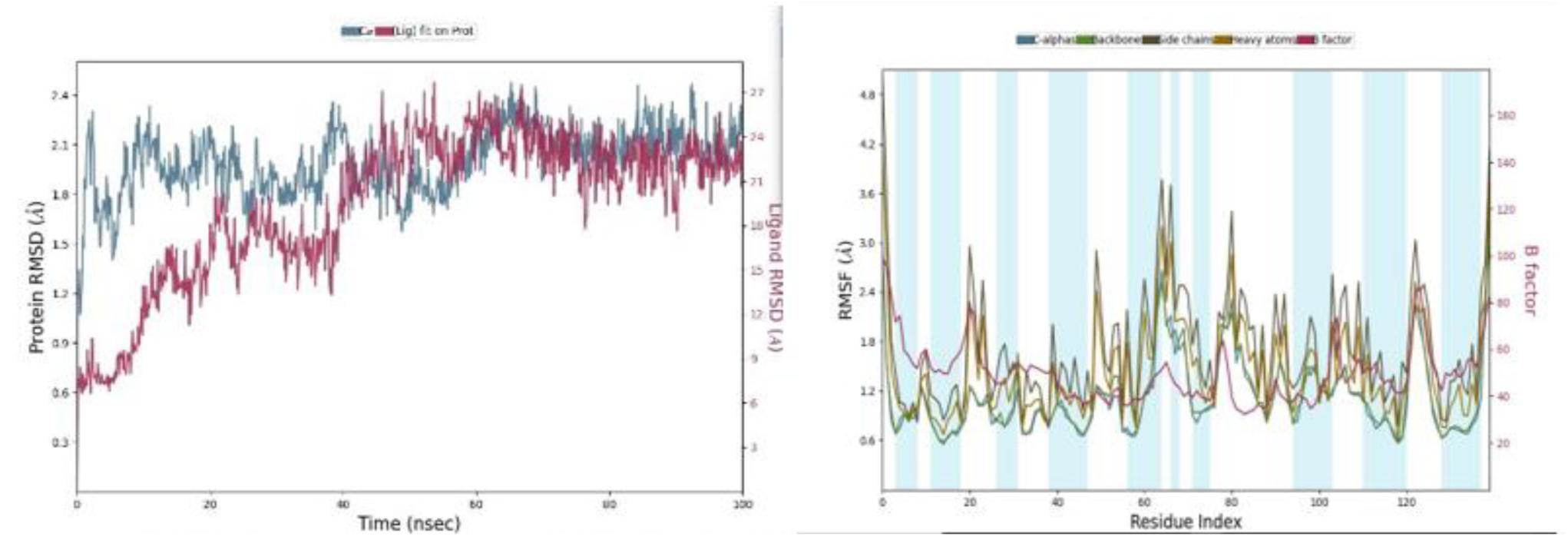
**In the RMSD plot (vaccine-receptor complex), blue peaks show Protein RMSD and red peaks indicate Ligand RMSD. Second image shows RMSF of TLR4 protein in the presence of constructed vaccine, as compared with B-factor.**

### Root mean square fluctuation (RMSF) calculations

A protein’s RMSF value is often determined to access side chain fluctuations caused due to ligand binding[71]. The larger RMSF values indicate flexibility during the simulation period, and the lower RMSF values show good system stability. In Figure 8, red and blue bars indicate the α-helices and β-sheets regions, while the loop area is white colour. The plot shows higher fluctuation in the C-terminal and N-terminal regions in contrast with other regions and minor fluctuations in RMSF values of TLR4 protein backbone and side chains in loop regions.

### Immunological Simulation

The immune response to the PKFDVac-I MEPV was evaluated using the C Imm-sim server through *in silico* simulations. The focus was on interactions of B-cell, class I&II HLA epitope and T-cell receptor with HLA-peptide complexes. Results showed that the vaccine triggered an immune response initially and even stronger responses in subsequent exposures. Higher levels of immunoglobulins and significant antigen reduction were observed post-vaccination (Figure 9).

**Figure 9:**
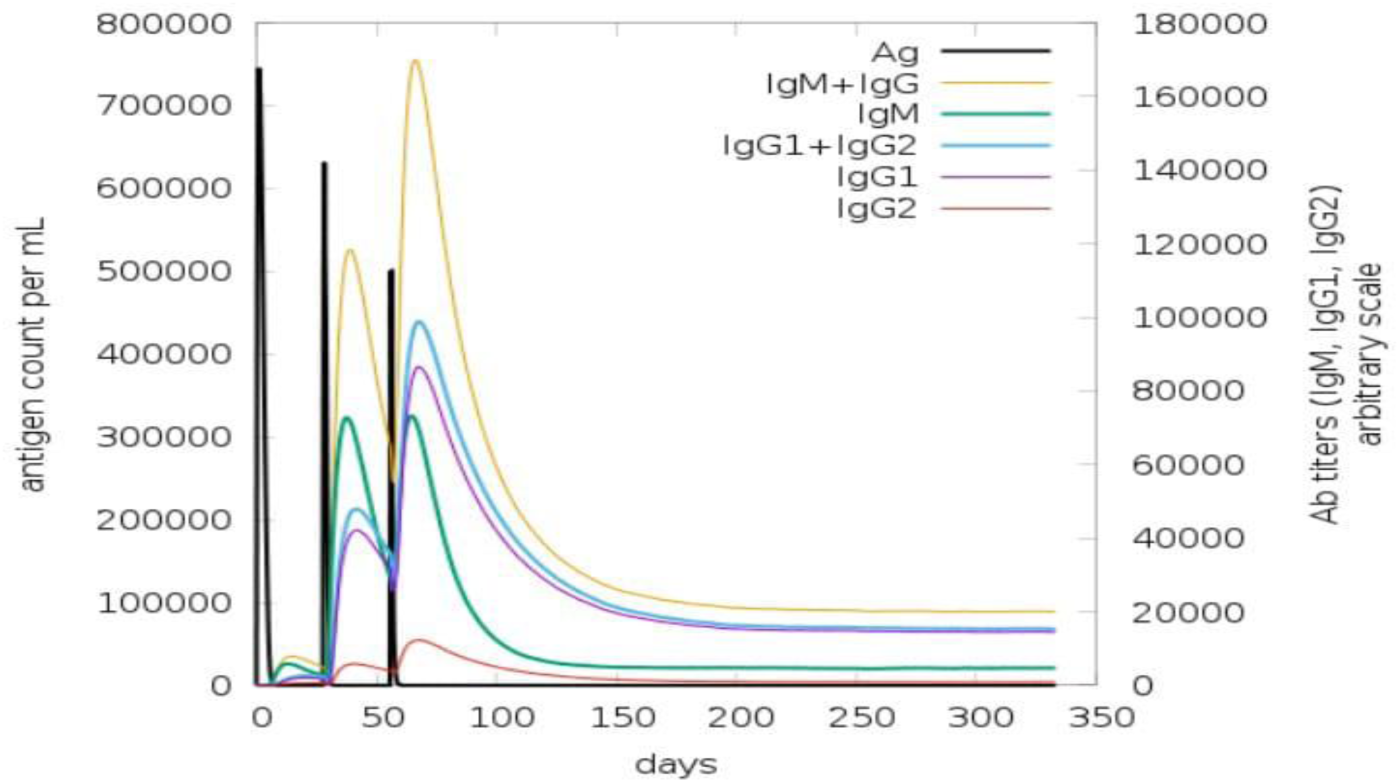
**Immune response developed against PKFDVac-I and the levels of immunoglobulins for the antigen concentration using the C-ImmSim server.**

The vaccine’s effectiveness was evidenced by the increased and sustained B-cell and T-cell populations, with each dose enhancing these levels and promoting the development of memory T-helper and T-cytotoxic cells. These cells remained active throughout the simulation, indicating a prolonged immune response. The T-cytotoxic cell population maintained steady memory cell levels and increased active cell levels. Additionally, the increased proliferation of macrophages and dendritic cells suggested efficient antigen processing and presentation to CD4+ and CD8+ cells. The simulation also showed elevated cytokine and interleukin levels, particularly a significant rise in Interferon Gamma, indicating a strong antiviral response generated by the PKFDVac-I vaccine construct.

## DISCUSSION

KFD popularly known as Monkey fever, found to have restricted distribution in Karnataka, India is recently found expanding its distribution along the stretch of Western Ghats. Annually, an estimated cases of around 400 KFD positives are documented yearly[72]. According to Kasabi et al[73] and Holbrook[74], the rate of KFD infected cases are increasing due to the lack of proper therapeutic methods and low efficacy of the existing vaccine. The conventional control measures such as the tick control and a CEF based vaccine was found to have limited effect over the cases. Since KFDV is regarded as a highly pathogenic agent and is categorized as BSL4[75] and KFDV was assigned a Risk Group 4 microbe classification as of 2017 in the document “Regulations and Guidelines on Biosafety of Recombinant DNA Research and Biocontainment”[4], using reverse vaccinology approach has made it simpler to identify the epitopes that trigger a potent immune response. To develop and formulate the MEPV construct, given the pathogenic characteristics of the virus, the existing genomic sequences of the pathogens were leveraged, and the multi-epitope antigenic peptides were predicted and the physiochemical characterstics were analysed through immunoinformatic tools listed in Table 2.

Among structural proteins, the KFD virus’ Envelope protein (E) is a significant tissue tropism cause. This protein is vital for the pathogenesis and immune evasion of the virus because it facilitates the virus’s entry into the host cell and determines its immunogenetic and phenotypic features[76]. A total of 25 HTL & CTL each and 66 B-cell epitopes were selected based on their binding score and percentile rank and were subjected to quality attributes analysis like antigenicity, allergenicity, toxicity, and immunogenicity.

After comprehensive analysis, 5 CTL, 3 HTL, and 8 B-cell epitopes which met the criteria for a viable vaccine candidate were finalized and linked using suitable linkers to create a stable and effective vaccine construct responsible potentially leading to a robust IR. The final PKFDVac-I vaccine comprised of 279 amino acids with an MW of 29,162.32 daltons, within the optimal size range for a vaccine. Physiochemical properties revealed an isoelectric point (pI) of 9.79, indicating the pH at which the protein is electrically neutral. The computed instability index (II) of 11.80. suggested protein stability, and the aliphatic index of 108.39 predicted thermal stability[65]. A GRAVY score of 0.448 indicated polarity and hydrophobicity, while a scaled solubility of 0.554 suggested high solubility[77].

Toll-like receptors (TLRs) form the first line of defense against infections and play a pivotal role in adaptive immunity. Specifically, TLRs 1-9, except TLR-5, are involved in responses to viral infections. However, few studies have shown that TLR4 can lead to deleterious immune responses in certain viral infections[78]. In our molecular docking experiments, the proposed vaccine PKFDVac-I candidate demonstrated high binding affinity and low binding energy (Piper energy −1150.78) at the TLR4 receptor binding site. Molecular dynamics simulation, a robust technique for analyzing the dynamics of protein-protein interactions, was used to evaluate the stability and mobility of the TLR4-vaccine complex within a biological-environment.

The PKFDVac-I, tested using computer simulations, elicited a strong immune response, particularly after repeated doses. The vaccine increased levels of key immune proteins and enhanced the activity of B-cells and T-cells, which are crucial for fighting infections. These immune cells remained active over an extended period. Additionally, the vaccine boosted the activity of antigen-presenting immune cells, which process and present antigens to other immune cells. Overall, the vaccine generated a potent antiviral response.

Given these promising results, PKFDVac-I warrants further studies through *in vitro* and *in vivo* studies to develop a safe and effective vaccine candidate against KFD for human use.

## CONCLUSION

KFD is a reemerging zoonotic problem in India and the resurgence of KFD in India highlights the necessity for effective preventive strategies that go beyond conventional vector management. The present study demonstrated the prediction of multiepitope peptide-based immunogens from the virus genome. The analysis has shown the possible application of these immunoinformatic predicted antigens as potential immunogens against KFD. From this study, the proposed PKFDVac-I vaccine demonstrated high binding affinity, stability, and promising immunogenicity *in silico*, implying that PKFDVac-I may function as a strong immunogen, indicating the need for more study to assess its effectiveness in suitable biological models. The work also emphasizes how immunoinformatically predicted antigens can also be chemically synthesised, which makes scaling up for large production simpler. Furthermore, it is worth determining any potential cross-protection provided by these immunogens *in vivo*, as this could expand the range of uses for them in the fight against related viral illnesses, especially considering the strong genetic resemblance between KFDV and the Alkhurma virus reported in Saudi Arabia.

## ACKNOWLEDGEMENT

The authors would like to thank Ms. Reshma M, Department of Microbiology, Pasteur Institute of India, Dr.Vijayakumar Rajendran, Indian Institute of Science, Banglore, India and Mr. Rajesh Nair for their continuous support and guidance.

## ABBREVIATIONS

AHFV: Alkhurma Hemorrhagic Fever Virus
BSL: Bio Safety Level
C Protein: Capsid Protein
CDC: Centers for Disease Control and Prevention
CEF: Chick Embryo Fibroblast
CFR: Case Fatality Rate
CTL: Cytotoxic T lymphocytes
E Protein: Envelope Protein
DMD: Discrete Molecular Dynamics
DNA: Deoxy Ribo Nucleic Acid
FASTA: Fast Adaptive Shrinkage Threshold Algorithm
GRAVY: Grand average of hydropathicity
HTL: Helper T lymphocytes
HLA: Human Leukocyte Antigen
IEDB: Immune Epitope Database
IFN: Interferon
II: Instability Index
IR: Immune Resposne
KFD: Kyasanur Forest Disease
KFDV: Kyasanur Forest Disease Virus
MD Simulation: Molecular Dynamics Simulation
MEPV: Multi Epitope Peptide Vaccine
MHC: Major Histocompatability Complex
MW: Molecular weight
NCBI: National Center for Biotechnology Information
NS Protein: Non Structural Protein
NPT: Constant number of particles, Pressure, and Temperature
PDB: Protein Data Bank
PKFDVac-I: PIIC Vac Candidate I
prM Protein: Precursor Membrane Protein
RMSF: Root Mean Square Fluctuation
RMSD: Root Mean Square Deviation
RNA: Ribo Nucleic Acid
RSSEV: Russian Spring-Sumer Encephalitis Virus
TLR: Toll like receptor

## AUTHOR CONTRIBUTIONS

Conceptualization and design: SBP, SJ; Methodology: SBP, AJ, SJ ; Computational work: SBP, AJ, NKP,RGP; Data analysis: SBP, SJ; Writing : SBP, SJ, PR ; Reviewing and Editing: SJB, PR ; Co-ordination of Research: SKC, AAP ; Supervision: SJ, SKC, PR

All authors read and approved the final manuscript.

## INFORMED CONSENT

Not applicable

## ETHICS STATEMENT

Not applicable

## REFERENCES

[1] T. H. Work, F. R. Roderiguez, and P. N. Bhatt, “Virological Epidemiology of the 1958 Epidemic of Kyasanur Forest Disease,” Am J Public Health Nations Health, vol. 49, no. 7, p. 869, 1959, doi: 10.2105/AJPH.49.7.869.

[2] N. K. Pariyapurath et al., “Targeted Immunization Strategies and Designing Vaccine against Indian Nipah Virus Strain (NiV B) and Malaysian Variant (NiV M),” Original Article International Journal of Pharmaceutical Investigation, vol. 14, no. 4, pp. 1201–1207, 2024, doi: 10.5530/ijpi.14.4.131.

[3] K. Pavri, “Clinical, Clinicopathologic, and Hematologic Features of Kyasanur Forest Disease,” Clinical Infectious Diseases, vol. 11, no. Supplement_4, pp. S854–S859, May 1989, doi: 10.1093/clinids/11.Supplement_4.S854.

[4] S. B. Pillai et al., “Monkeypox and Monkey Fever: Basic Understanding for Better Community Participation in Disease Control,” Biotechnology Journal International, vol. 26, no. 5, pp. 1–12, Oct. 2022, doi: 10.9734/bji/2022/v26i5657.

[5] D. Y. Patil et al., “Occupational exposure of cashew nut workers to Kyasanur Forest disease in Goa, India,” International Journal of Infectious Diseases, vol. 61, pp. 67–69, Aug. 2017, doi: 10.1016/J.IJID.2017.06.004.

[6] S. Chakraborty, W. Sander, B. F. Allan, and F. C. D. Andrade, “Sociodemographic factors associated with Kyasanur forest disease in India - a retrospective study.,” IJID regions, vol. 10, pp. 219–227, Mar. 2024, doi: 10.1016/j.ijregi.2024.02.002.

[7] A. P. Sebastian, M. Varma, and N. Gupta, “Kyasanur forest disease in India: a case report.,” J Travel Med, vol. 31, no. 5, Jul. 2024, doi: 10.1093/jtm/taae071.

[8] “Kyasanur Forest Disease: A public health concern, CD Alert,” National Centre for Disease Control, Delhi. Accessed: Sep. 17, 2022. [Online]. Available: https://ncdc.mohfw.gov.in/wp-content/uploads/2024/05/KYASANUR.pdf

[9] K. Chunduru and K. Saravu, “Kyasanur forest disease: A review on the emerging infectious disease,” Journal of Clinical Infectious Diseases Society, vol. 1, no. 1, p. 5, 2023, doi: 10.4103/cids.cids_13_23.

[10] J. Wang et al., “Isolation of kyasanur forest disease virus from febrile patient, Yunnan, China,” Emerg Infect Dis, vol. 15, no. 2, pp. 326–328, Feb. 2009, doi: 10.3201/eid1502.080979.

[11] B. Bhatia, H. Feldmann, and A. Marzi, “Kyasanur Forest Disease and Alkhurma Hemorrhagic Fever Virus-Two Neglected Zoonotic Pathogens.,” Microorganisms, vol. 8, no. 9, pp. 1–17, Sep. 2020, doi: 10.3390/microorganisms8091406.

[12] K. A. Dodd, B. H. Bird, M. E. B. Jones, S. T. Nichol, and C. F. Spiropoulou, “Kyasanur forest disease virus infection in mice is associated with higher morbidity and mortality than infection with the closely related Alkhurma hemorrhagic fever virus,” PLoS One, vol. 9, no. 6, p. e10031, Jun. 2014, doi: 10.1371/journal.pone.0100301.

[13] N. Gupta, W. Wilson, A. Neumayr, and K. Saravu, “Kyasanur forest disease: a state-of-the-art review.,” QJM, vol. 115, no. 6, pp. 351–358, Jun. 2022, doi: 10.1093/qjmed/hcaa310.

[14] A. Munivenkatappa, R. Sahay, P. Yadav, R. Viswanathan, and D. Mourya, “Clinical & epidemiological significance of Kyasanur forest disease,” Indian Journal of Medical Research, vol. 148, no. 2, p. 145, 2018, doi: 10.4103/ijmr.IJMR_688_17.

[15] M. V Murhekar, G. S. Kasabi, S. M. Mehendale, D. T. Mourya, P. D. Yadav, and B. V Tandale, “On the transmission pattern of Kyasanur Forest disease (KFD) in India.,” Infect Dis Poverty, vol. 4, p. 37, Aug. 2015, doi: 10.1186/s40249-015-0066-9.

[16] B. Bhatia, K. Meade-White, E. Haddock, F. Feldmann, A. Marzi, and H. Feldmann, “A live-attenuated viral vector vaccine protects mice against lethal challenge with Kyasanur Forest disease virus,” NPJ Vaccines, vol. 6, no. 1, Dec. 2021, doi: 10.1038/s41541-021-00416-2.

[17] G. S. Kasabi et al., “Coverage and Effectiveness of Kyasanur Forest Disease (KFD) Vaccine in Karnataka, South India, 2005-10,” PLoS Negl Trop Dis, vol. 7, no. 1, p. e2025, 2013, doi: 10.1371/journal.pntd.0002025.

[18] S. K. Kiran et al., “Kyasanur Forest disease outbreak and vaccination strategy,Shimoga District, India, 2013-2014,” Emerg Infect Dis, vol. 21, no. 1, pp. 146–149, Jan. 2015, doi: 10.3201/eid2101.141227.

[19] C. B. Jonsson et al., “Epidemiology, Pathogenesis, and Control of a Tick-Borne Disease-Kyasanur Forest Disease: Current Status and Future Directions,” Frontiers in Cellular and Infection Microbiology | www.frontiersin.org, vol. 1, p. 149, 2018, doi: 10.3389/fcimb.2018.00149.

[20] P. Rajaiah, “Kyasanur Forest Disease in India: innovative options for intervention.,” Hum Vaccin Immunother, vol. 15, no. 10, pp. 2243–2248, 2019, doi: 10.1080/21645515.2019.1602431.

[21] K. A. Dodd et al., “Ancient Ancestry of KFDV and AHFV Revealed by Complete Genome Analyses of Viruses Isolated from Ticks and Mammalian Hosts,” PLoS Negl Trop Dis, vol. 5, no. 10, p. e1352, Oct. 2011, doi: 10.1371/journal.pntd.0001352.

[22] X. Zhang, Y. Zhang, R. Jia, M. Wang, Z. Yin, and A. Cheng, “Structure and function of capsid protein in flavivirus infection and its applications in the development of vaccines and therapeutics,” Vet Res, vol. 52, no. 1, p. 98, Jun. 2021, doi: 10.1186/s13567-021-00966-2.

[23] D. T. Mourya et al., “Diagnosis of Kyasanur forest disease by nested RT-PCR, real-time RT-PCR and IgM capture ELISA,” J Virol Methods, vol. 186, no. 1–2, pp. 49–54, Dec. 2012, doi: 10.1016/J.JVIROMET.2012.07.019.

[24] D. Lin et al., “Analysis of the complete genome of the tick-borne flavivirus Omsk hemorrhagic fever virus,” Virology, vol. 313, no. 1, pp. 81–90, Aug. 2003, doi: 10.1016/S0042-6822(03)00246-0.

[25] P. Shil, P. D. Yadav, A. A. Patil, R. Balasubramanian, and D. T. Mourya, “Bioinformatics characterization of envelope glycoprotein from kyasanur forest disease virus,” Indian Journal of Medical Research, vol. 147, no. February, pp. 195–201, Feb. 2018, doi: 10.4103/ijmr.IJMR_1445_16.

[26] B. S. Pillai, S. Jagannathan, S. Jagadibabu, C. S. Kukkaler, and S. Sakthivel, “Immunodominant Protein of Kyasanur Forest Disease Virus: A retrospective study,” Res J Biotechnol, vol. 19, no. 9, pp. 132–142, Jul. 2024, doi: 10.25303/1909rjbt1320142.

[27] S. Dey, M. Pratibha, H. Singh Dagur, and E. Rajakumara, “Characterization of host receptor interaction with envelop protein of Kyasanur forest disease virus and predicting suitable epitopes for vaccine candidate.,” J Biomol Struct Dyn, vol. 42, no. 8, pp. 4110–4120, May 2024, doi: 10.1080/07391102.2023.2218924.

[28] S. Odend’hal, “Kyasanur Forest Disease Virus,” The Geographical Distribution of Animal Viral Diseases, vol. 1, pp. 253–256, 1983, doi: 10.1016/B978-0-12-524180-9.50070-4.

[29] I. A. Doytchinova and D. R. Flower, “VaxiJen: A server for prediction of protective antigens, tumour antigens and subunit vaccines,” BMC Bioinformatics, vol. 8, Jan. 2007, doi: 10.1186/1471-2105-8-4.

[30] F. F. Gonzalez-Galarza et al., “Allele frequency net database (AFND) 2020 update: Gold-standard data classification, open access genotype data and new query tools,” Nucleic Acids Res, vol. 48, no. D1, pp. D783–D788, Jan. 2020, doi: 10.1093/nar/gkz1029.

[31] A. Jeyachandran et al., “Identification and evaluation of multi-antigenic epitopes of immunodominant protein from the selected Crimean–Congo hemorrhagic fever virus genome towards the development of diagnostic and vaccine candidates by reverse vaccinology approach,” J Proteins Proteom, Sep. 2024, doi: 10.1007/s42485-024-00164-6.

[32] B. Reynisson, B. Alvarez, S. Paul, B. Peters, and M. Nielsen, “NetMHCpan-4.1 and NetMHCIIpan-4.0: improved predictions of MHC antigen presentation by concurrent motif deconvolution and integration of MS MHC eluted ligand data.,” Nucleic Acids Res, vol. 48, no. W1, pp. W449–W454, Jul. 2020, doi: 10.1093/nar/gkaa379.

[33] S. Saha and G. P. S. Raghava, “Prediction of continuous B-cell epitopes in an antigen using recurrent neural network.,” Proteins, vol. 65, no. 1, pp. 40–8, Oct. 2006, doi: 10.1002/prot.21078.

[34] J. Ponomarenko et al., “ElliPro: a new structure-based tool for the prediction of antibody epitopes,” 2008, doi: 10.1186/1471-2105-9-514.

[35] J. Erik, P. Larsen, O. Lund, and M. Nielsen, “Improved method for predicting linear B-cell epitopes,” 2006, doi: 10.1186/1745-7580-2-2.

[36] S. Saha and G. P. S. Raghava, “Prediction methods for B-cell epitopes.,” Methods Mol Biol, vol. 409, pp. 387–94, 2007, doi: 10.1007/978-1-60327-118-9_29.

[37] M. B. Stadler and B. M. Stadler, “Allergenicity prediction by protein sequence.,” FASEB J, vol. 17, no. 9, pp. 1141–3, Jun. 2003, doi: 10.1096/fj.02-1052fje.

[38] I. Dimitrov, I. Bangov, D. R. Flower, and I. Doytchinova, “AllerTOP v.2 - A server for in silico prediction of allergens,” J Mol Model, vol. 20, no. 6, 2014, doi: 10.1007/s00894-014-2278-5.

[39] N. Sharma, S. Patiyal, A. Dhall, A. Pande, C. Arora, and G. P. S. Raghava, “AlgPred 2.0: an improved method for predicting allergenic proteins and mapping of IgE epitopes.,” Brief Bioinform, vol. 22, no. 4, Jul. 2021, doi: 10.1093/bib/bbaa294.

[40] S. Gupta et al., “In silico approach for predicting toxicity of peptides and proteins.,” PLoS One, vol. 8, no. 9, p. e73957, 2013, doi: 10.1371/journal.pone.0073957.

[41] S. K. Dhanda et al., “Predicting HLA CD4 Immunogenicity in Human Populations,” Front Immunol, vol. 9, p. 1369, Jun. 2018, doi: 10.3389/fimmu.2018.01369.

[42] S. K. Dhanda, P. Vir, and G. P. S. Raghava, “Designing of interferon-gamma inducing MHC class-II binders.,” Biol Direct, vol. 8, p. 30, Dec. 2013, doi: 10.1186/1745-6150-8-30.

[43] H.-H. Bui, J. Sidney, W. Li, N. Fusseder, and A. Sette, “Development of an epitope conservancy analysis tool to facilitate the design of epitope-based diagnostics and vaccines.,” BMC Bioinformatics, vol. 8, p. 361, Sep. 2007, doi: 10.1186/1471-2105-8-361.

[44] J. Dey et al., “Designing a novel multi-epitope vaccine to evoke a robust immune response against pathogenic multidrug-resistant Enterococcus faecium bacterium,” Gut Pathog, vol. 14, no. 1, p. 21, May 2022, doi: 10.1186/s13099-022-00495-z.

[45] Z. Nain et al., “Proteome-wide screening for designing a multi-epitope vaccine against emerging pathogen Elizabethkingia anophelis using immunoinformatic approaches.,” J Biomol Struct Dyn, vol. 38, no. 16, pp. 4850–4867, Oct. 2020, doi: 10.1080/07391102.2019.1692072.

[46] S. I. Islam, S. Mahfuj, Md. A. Alam, Y. Ara, S. Sanjida, and M. J. Mou, “Immunoinformatic Approaches to Identify Immune Epitopes and Design an Epitope-Based Subunit Vaccine against Emerging Tilapia Lake Virus (TiLV),” Aquaculture Journal, vol. 2, no. 2, pp. 186–202, Jun. 2022, doi: 10.3390/aquacj2020010.

[47] E. Gasteiger et al., “Protein Identification and Analysis Tools on the ExPASy Server,” in *The Proteomics Protocols Handbook*, Totowa, NJ: Humana Press, 2005, pp. 571–607. doi: 10.1385/1-59259-890-0:571.

[48] M. Hebditch, M. A. Carballo-Amador, S. Charonis, R. Curtis, and J. Warwicker, “Protein-Sol: a web tool for predicting protein solubility from sequence.,” Bioinformatics, vol. 33, no. 19, pp. 3098–3100, Oct. 2017, doi: 10.1093/bioinformatics/btx345.

[49] D. T. Jones, “Protein secondary structure prediction based on position-specific scoring matrices.,” J Mol Biol, vol. 292, no. 2, pp. 195–202, Sep. 1999, doi: 10.1006/jmbi.1999.3091.

[50] C. Geourjon and G. Deléage, “SOPMA: significant improvements in protein secondary structure prediction by consensus prediction from multiple alignments,” Bioinformatics, vol. 11, no. 6, pp. 681–684, 1995, doi: 10.1093/bioinformatics/11.6.681.

[51] J. Jumper et al., “Highly accurate protein structure prediction with AlphaFold,” Nature, vol. 596, no. 7873, pp. 583–589, Aug. 2021, doi: 10.1038/s41586-021-03819-2.

[52] R. A. Laskowski, “Enhancing the functional annotation of PDB structures in PDBsum using key figures extracted from the literature,” Bioinformatics, vol. 23, no. 14, pp. 1824–1827, Jul. 2007, doi: 10.1093/bioinformatics/btm085.

[53] S. Ramachandran, P. Kota, F. Ding, and N. V Dokholyan, “Automated minimization of steric clashes in protein structures.,” Proteins, vol. 79, no. 1, pp. 261–70, Jan. 2011, doi: 10.1002/prot.22879.

[54] Maestro, “Schrödinger Release 2024-3.”

[55] T. A. Bouback et al., “Pharmacophore-based virtual screening, quantum mechanics calculations, and molecular dynamics simulation approaches identified potential natural antiviral drug candidates against MERS-CoV S1-NTD,” Molecules, vol. 26, no. 16, Aug. 2021, doi: 10.3390/molecules26164961.

[56] N. Rapin, O. Lund, M. Bernaschi, and F. Castiglione, “Computational immunology meets bioinformatics: The use of prediction tools for molecular binding in the simulation of the immune system,” PLoS One, vol. 5, no. 4, 2010, doi: 10.1371/journal.pone.0009862.

[57] F. Castiglione, F. Mantile, P. De Berardinis, and A. Prisco, “How the interval between prime and boost injection affects the immune response in a computational model of the immune system.,” Comput Math Methods Med, vol. 2012, p. 842329, 2012, doi: 10.1155/2012/842329.

[58] A. K. Rouzbahani, F. Kheirandish, and S. Z. Hosseini, “Design of a multi-epitope-based peptide vaccine against the S and N proteins of SARS-COV-2 using immunoinformatics approach.,” The Egyptian journal of medical human genetics, vol. 23, no. 1, p. 16, 2022, doi: 10.1186/s43042-022-00224-w.

[59] A. Naz, F. Shahid, T. T. Butt, F. M. Awan, A. Ali, and A. Malik, “Designing Multi-Epitope Vaccines to Combat Emerging Coronavirus Disease 2019 (COVID-19) by Employing Immuno-Informatics Approach.,” Front Immunol, vol. 11, p. 1663, 2020, doi: 10.3389/fimmu.2020.01663.

[60] K. Venugopal, T. Gritsun, V. A. Lashkevich, and E. A. Gould, “Analysis of the structural protein gene sequence shows Kyasanur Forest disease virus as a distinct member in the tick-borne encephalitis virus serocomplex.,” J Gen Virol, vol. 75 ( Pt 1), pp. 227–32, Jan. 1994, doi: 10.1099/0022-1317-75-1-227.

[61] P. Shil, P. D. Yadav, A. A. Patil, R. Balasubramanian, and D. T. Mourya, “Bioinformatics characterization of envelope glycoprotein from kyasanur forest disease virus,” Indian Journal of Medical Research, vol. 147, no. February, pp. 195–201, Feb. 2018, doi: 10.4103/ijmr.IJMR_1445_16.

[62] D.S.Burke and T.P.Monath, *Fields Virology*, 4th Edition. Philadelphia: Lippincott Williams and Wilkins, 2001.

[63] H. Holzmann, F. X. Heinz, C. W. Mandl, F. Guirakhoo, and C. Kunz, “A single amino acid substitution in envelope protein E of tick-borne encephalitis virus leads to attenuation in the mouse model.,” J Virol, vol. 64, no. 10, pp. 5156–5159, Oct. 1990, doi: 10.1128/JVI.64.10.5156-5159.1990.

[64] S. Baseer, S. Ahmad, K. E. Ranaghan, and S. S. Azam, “Towards a peptide-based vaccine against Shigella sonnei: A subtractive reverse vaccinology based approach.,” Biologicals, vol. 50, pp. 87–99, Nov. 2017, doi: 10.1016/j.biologicals.2017.08.004.

[65] D. D. Martinelli, “In silico vaccine design: A tutorial in immunoinformatics,” Healthcare Analytics, vol. 2, p. 100044, Nov. 2022, doi: 10.1016/j.health.2022.100044.

[66] O. Mejias-Gomez et al., “A window into the human immune system: comprehensive characterization of the complexity of antibody complementary-determining regions in functional antibodies,” MAbs, vol. 15, no. 1, 2023, doi: 10.1080/19420862.2023.2268255.

[67] D. W. A. Buchan and D. T. Jones, “The PSIPRED Protein Analysis Workbench: 20 years on.,” Nucleic Acids Res, vol. 47, no. W1, pp. W402–W407, Jul. 2019, doi: 10.1093/nar/gkz297.

[68] M. Mirdita, K. Schütze, Y. Moriwaki, L. Heo, S. Ovchinnikov, and M. Steinegger, “ColabFold: making protein folding accessible to all,” Nat Methods, vol. 19, no. 6, pp. 679–682, Jun. 2022, doi: 10.1038/s41592-022-01488-1.

[69] “The docking algorithms.” Accessed: Sep. 17, 2023. [Online]. Available: https://resources.qiagenbioinformatics.com/manuals/clcdrugdiscoveryworkbench/200/_docking_algorithms.html#:~:text=The%20score%20mimics%20the%20potential,weak%20or%20non%2Dexisting%20binding.

[70] S. R. Acharyya, P. Sen, T. Kandasamy, and S. S. Ghosh, “Designing of disruptor molecules to restrain the protein-protein interaction network of VANG1/SCRIB/NOS1AP using fragment-based drug discovery techniques.,” Mol Divers, vol. 27, no. 3, pp. 989–1010, Jun. 2023, doi: 10.1007/s11030-022-10462-0.

[71] G. Muteeb, A. Alshoaibi, M. Aatif, M. T. Rehman, and M. Z. Qayyum, “Screening marine algae metabolites as high-affinity inhibitors of SARS-CoV-2 main protease (3CLpro): an in silico analysis to identify novel drug candidates to combat COVID-19 pandemic.,” Appl Biol Chem, vol. 63, no. 1, p. 79, 2020, doi: 10.1186/s13765-020-00564-4.

[72] S. Z. Shah et al., “Epidemiology, Pathogenesis, and Control of a Tick-Borne Disease-Kyasanur Forest Disease: Current Status and Future Directions,” Front Cell Infect Microbiol, vol. 8, p. 149, May 2018, doi: 10.3389/fcimb.2018.00149.

[73] G. S. Kasabi et al., “Coverage and Effectiveness of Kyasanur Forest Disease (KFD) Vaccine in Karnataka, South India, 2005-10,” PLoS Negl Trop Dis, vol. 7, no. 1, 2013, doi: 10.1371/journal.pntd.0002025.

[74] M. R. Holbrook, “Kyasanur forest disease.,” Antiviral Res, vol. 96, no. 3, pp. 353–62, Dec. 2012, doi: 10.1016/j.antiviral.2012.10.005.

[75] M. Muraleedharan, “Kyasanur Forest Disease (KFD): Rare Disease of Zoonotic Origin.,” J Nepal Health Res Counc, vol. 14, no. 34, pp. 214–218, Sep. 2016.

[76] W. C. Barker, R. Mazumder, S. Vasudevan, J.-L. Sagripanti, and C. H. Wu, “Sequence signatures in envelope protein may determine whether flaviviruses produce hemorrhagic or encephalitic syndromes,” Virus Genes, vol. 39, no. 1, pp. 1–9, Aug. 2009, doi: 10.1007/s11262-009-0343-4.

[77] E. Gasteiger et al., “Protein Identification and Analysis Tools on the ExPASy Server,” in *The Proteomics Protocols Handbook*, Totowa, NJ: Humana Press, 2005, pp. 571–607. doi: 10.1385/1-59259-890-0:571.

[78] M. Carty and A. G. Bowie, “Recent insights into the role of Toll-like receptors in viral infection.,” Clin Exp Immunol, vol. 161, no. 3, pp. 397–406, Sep. 2010, doi: 10.1111/j.1365-2249.2010.04196.x.

